# Vascular signals coordinate cerebellar circuitry development

**DOI:** 10.64898/2026.02.18.706533

**Authors:** M. Parrilla, R. Hülse, J. Jin, A. D’Errico, J. Redondo-Nectalí, S. Srivastava, F. Stuth, N. Alivodej, C. Llaó-Cid, M.R. Aburto, M. Segarra, A. Acker-Palmer

**Author notes:** Equal contribution (R. Hülse, J. Jin).

## Abstract

Neural circuit assembly requires precise coordinated interactions between developing neurons and the vasculature, yet the instructive signals provided by endothelial cells remain largely unknown. Here, we identify a vascular-to-neural signaling axis that orchestrates postnatal cerebellar development. Endothelial-specific deletion of the adaptor protein Dab1 in mice disrupted vascular patterning and uncoupled the growth of major cerebellar neuronal populations. We show that endothelial Dab1 in the cerebellum drives secretion of the morphogen Wnt5a, which acts through Frizzled-2 to restrain granule-cell progenitor proliferation and promote Purkinje-cell dendritic maturation. Endothelial Wnt5a deletion phenocopied Dab1 endothelial mutant (*Dab1*^iΔEC^) defects, whereas exogenous Wnt5a restored normal progenitor dynamics in *Dab1*^iΔEC^ cerebellar slices, demonstrating pathway sufficiency. Functionally, loss of this vascular signal impaired Purkinje-cell firing, diminished parallel-fiber input, reduced synapse formation from both parallel and climbing fibers, and disrupted long-term plasticity. These findings reveal a key instructive role for blood vessels in shaping cerebellar architecture and establishing functional circuit connectivity.

## Introduction

The cerebellum, long recognized for its role in motor coordination, is now appreciated as a hub for cognitive and social processing ^1,2^. Its diverse functions rely on the precise assembly of neural circuits during development, beginning in embryogenesis with the spatial and temporal specification of neuronal progenitors ^3^. While key developmental steps occur in utero, a substantial portion of cerebellar maturation, including neuronal proliferation, migration, and synaptogenesis, takes place postnatally in both mice and humans ^3^.

Granule cells (GCs), the most abundant neurons in the brain, are excitatory glutamatergic interneurons that synapse onto Purkinje cells (PCs), GABAergic inhibitory neurons that constitute the sole output of the cerebellar cortex ^3–5^. Granule cell progenitors (GCP) proliferate in the outer external granule layer (EGL), a transient germinal zone located just beneath the pial vasculature, and subsequently migrate inward to form the internal granule layer (GCL), leaving behind parallel fibers (PF) that establish synapses with the nascent PC dendrites ^3–5^. Concurrently, PCs arrange their somata in a monolayer, known as Purkinje cell layer (PCL) and elaborate their extensive dendritic arbors into the nascent molecular layer (ML) ^3,6,7^. By the third to fourth postnatal week, the cerebellar cortex reaches its mature laminar cytoarchitecture and its synaptic organization ^3,5^.

An emerging body of work suggests that the vasculature actively shapes neural development beyond its canonical function of oxygen and nutrient delivery (reviewed in ^8^). Vessels can modulate neurogenesis ^9^ and guide neuronal migration in multiple brain regions ^10,11^. In the cerebellum, changes in vascular density and oxygen tension have been shown to influence GCP proliferation ^12^. However, beyond regulating tissue oxygenation, the instructive roles of endothelial cells in postnatal cerebellar development, particularly in orchestrating circuit assembly and neuronal maturation, remain largely unexplored.

Here, we identify an endothelial-to-neuronal signaling axis that is essential for the coordinated development of cerebellar circuits. Using temporally controlled, endothelium-specific genetic deletions, we show that endothelial Dab1 signaling governs both GCP proliferation and PC maturation. Dab1 controls the secretion of the non-canonical Wnt ligand Wnt5a from blood vessels, and that Wnt5a signals through the Frizzled-2 (Fzd2) receptor expressed on both GCPs and PCs. Pial and sub-PCL vasculature emerge as key sources of Wnt5a during the critical window of cerebellar circuit assembly. Loss of Wnt5a in endothelial cells recapitulates the vascular Dab1 mutant, whereas exogenous Wnt5a restores normal GPC proliferative dynamics. Furthermore, the loss of endothelial Dab1 produces long-lasting structural and functional defects that consist of misaligned PF axons, reduced excitatory synaptic input from both PFs and climbing fibers, impaired PC firing, and defective long-term synaptic plasticity.

Together, these findings establish the vasculature not merely as a passive scaffold or metabolic support, but as an active instructive player in postnatal cerebellar development. By defining a vascular-to-neural Dab1–Wnt5a axis that couples vascular cues to neuronal proliferation, dendritic maturation, and synaptic integration, our study provides a mechanistic framework for how endothelial cells orchestrate the temporal coordination of cerebellar circuit assembly.

## Results

### Postnatal sprouting angiogenesis parallels cerebellar neuronal development

The development of the cerebellum during the early postnatal period is marked by rapid changes in both neuronal and vascular architecture. While it is well established that cerebellar neurons such as GCs and PCs mature extensively after birth, it remains unclear how vascular growth unfolds in parallel and whether this spatially organized angiogenesis is coordinated with neuronal development. To address this, we characterized the temporal and spatial dynamics of cerebellar blood vessels from birth through the second postnatal week. At postnatal day 1 (P1), we observed initial vascular sprouts extending from the pial surface into the cerebellar cortex. These sprouts reached the incipient PCL, where Calbindin-positive (Cb⁺) PCs had not yet formed a monolayer (Figure 1A). Vascular branching in the EGL was minimal at this stage. A similar pattern was observed at P4, with pial vessels continuing to penetrate the cortex while the deeper layers remained largely devoid of branches (Figure 1B).

**Fig. 1.**
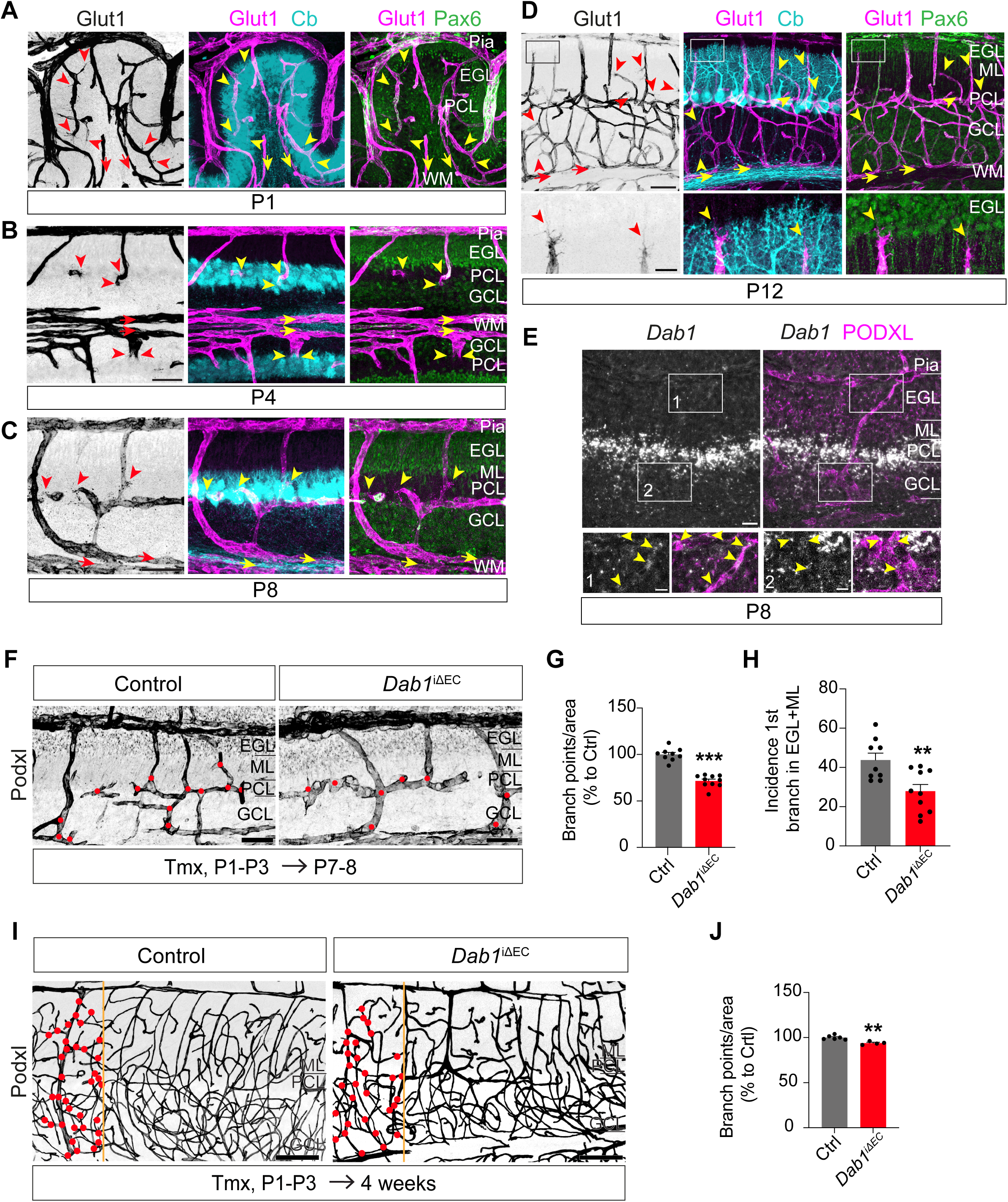
Postnatal cerebellar angiogenesis parallels neuronal layer formation and requires endothelial Dab1. (**A-D**) Immunolabeling of blood vessels with anti-Glut1 (black or magenta), Purkinje cells with anti-Calbindin (Cb, cyan) and granule cells with anti-Pax6 (green) at P1 (A), P4 (B), P8 (C) and P12 (D) from C57BL/6 mice. Arrowheads indicate vascular sprouting in the cerebellar cortex. Arrows indicate vessels in the white matter. Higher magnification of D below showing sprouting vessels in the molecular layer directed to the external granule layer. (**E**) *Dab1* mRNA detected by fluorescent *in situ* hybridization in cerebellar cortex at P8. Blood vessels were immunostained for the vessel marker Podocalyxin (Podxl). Higher magnifications are shown below. Arrows indicate *Dab1* signal in the vasculature. (**F**) *Dab1*^iΔEC^ cerebellar cortical vessels stained with Podxl at P8 after tamoxifen (Tmx) administration from P1 to P3. Red dots indicate the branch points. (**G-H**) Quantification of branch point per area (G) and distribution of incidence of the first branch point in both layers EGL and ML (H). Vascularization in the mutant cerebellar cortices is reduced, showing fewer branching points per area and less frequency of finding the first branch point in the EGL+ML (n = 9-10 animals per genotype). (**I**) Blood vessels were immunolabelled with anti-Podxl in *Dab1*^iΔEC^ and control cerebellar cortices in 4-week-old mice after Tmx administration from P1 to P3. Red dots in the region limited by the orange line indicate the branch points. (**J**) Quantification of branch points per area. Vascularization in the mutant cerebellar cortices is reduced showing fewer branching points. (n = 4-6 animals per genotype). Abbreviations: EGL = external granule layer, ML = molecular layer, PCL = Purkinje cell layer, GCL = granule cell layer and WM = white matter. Scale bars: A-D = 50 µm; higher magnification in D and E = 10 µm; E = 20 µm; F = 50 µm; I = 100 µm. Data are shown as mean ± SEM. \*\**P*<0.01, \*\*\**P*<0.001.

By P8, cerebellar neurons had adopted more defined spatial arrangements. PCs were organized into a clear monolayer within the PCL, and the vasculature showed increased complexity. Blood vessels aligned longitudinally beneath the PC somas and exhibited active sprouting into the ML, coinciding with the early stages of PC dendritogenesis and GC synaptogenesis (Figure 1C). At the same time, vascular branching within the GCL was notably increased compared to earlier stages.

At P12, we detected extensive vessel sprouting and branching throughout both the ML and GCL. In contrast, the EGL remained poorly vascularized, indicating that vessels entering in the EGL underwent little to no local sprouting (Figure 1D). Intriguingly, some vessels originating in the PCL or ML re-colonized the EGL and extended filopodia back toward the pial surface, suggesting the existence of anastomotic connections forming between deep and superficial vascular networks (Figure 1D).

Throughout this early postnatal window, white matter vessels maintain a distinctive arrangement, closely surrounding the extending PC axons (Figure 1A-D), raising the possibility that specific vessel–axon interactions may also play instructive roles in organizing long-range cerebellar projections.

Altogether, these findings revealed a highly structured and dynamic angiogenic process that unfolded in parallel with neuronal maturation. The timing and spatial distribution of vascular growth appeared to mirror the emergence of cerebellar lamination and circuitry, suggesting coordinated development of vascular and neuronal architectures.

### Endothelial Dab1 regulates postnatal cerebellar vascular branching

Reelin is an essential neuronal guidance cue for all layered structures of the CNS, including the cerebellum ^13–15^. In agreement with previous reports ^16^, we detected Reelin protein in the EGL and GCL and was predominantly expressed by granule cells in the inner EGL, suggesting that postmitotic granule cell progenitors (GCPs) are its primary source (Figure S1A). Fluorescent in situ hybridization at P8 revealed that Dab1 was not only expressed in PCs, as previously described ^17^, but was also present in the pial vasculature, in penetrating vessels within the EGL and in the vascular plexus located beneath the PCL (Figure 1E).

To dissect the function of Dab1 in endothelial cells, we generated an inducible endothelial-specific Dab1 knockout mouse line (*Dab1*^iΔEC^) by crossing *Dab1^lox/lox^* mice with *Cdh5-creERT2* mice ^10^. 4OH-Tamoxifen was administered from P1 to P3, and cerebellar cortices were analyzed at P8. Immunolabeling for the vessel marker Podocalyxin revealed that blood vessels in control mice exhibited extensive branching beneath the PCL and in the GCL, with additional sparse sprouting into the nascent ML (Figure 1F). In contrast, *Dab1*^iΔEC^ mice displayed a marked reduction in vascular complexity. Quantification showed approximately a 30% decrease in branch points per area compared to controls (Figure 1F and 1G). The incidence of the first branch point within the EGL and ML was also significantly lower in *Dab1*^iΔEC^ mice (Figure 1F and 1H). Vascular branching remained reduced at four weeks of age (Figure 1I and 1J).

### Endothelial Dab1 restricts granule cell progenitor proliferation

We observed that the vascular branching phenotype in *Dab1*^iΔEC^ mice was particularly evident in proximity to the proliferative zone of the EGL and beneath the PCL. Given this spatial overlap, we asked whether the absence of endothelial Dab1 could also influence the behavior of GCPs, which proliferate near the pial surface during early postnatal development. During this period, GCPs in the EGL proliferate rapidly, with mitotic activity peaking around P8 ^18^. Using phospho-Histone H3 (pH3) as a marker for M-phase cells, we observed a significant increase in proliferating GCPs in *Dab1*^iΔEC^ mice compared to controls (Figure 2A and 2B). This phenotype was confirmed using Ki67 immunostaining, which labels cells in all active phases of the cell cycle and revealed an increased population of cycling GCPs (Figure S2A and S2B). Conversely, expression of the postmitotic marker p27 Kip1 was reduced in *Dab1*^iΔEC^ cerebella, indicating a disruption in GCP cell cycle exit (Figure 2A and 2C).

**Fig. 2.**
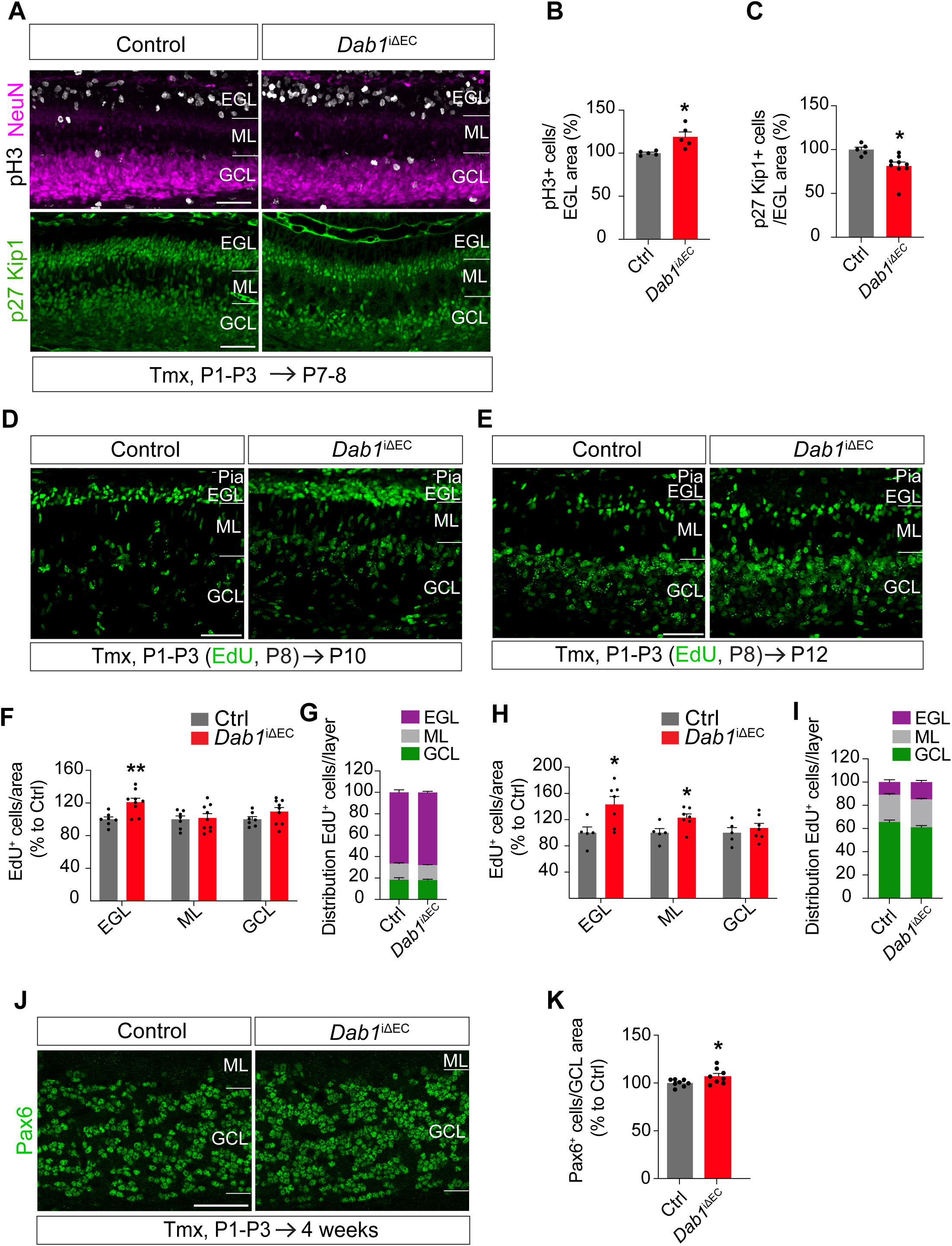
Endothelial Dab1 restrains granule cell progenitor proliferation during postnatal cerebellar development. (**A**) anti-pH3 and anti-p27 Kip1 immunostaining was used to determine the cell cycle stage of granule cell precursors in *Dab1*^iΔEC^ cerebellar cortices at P8 after tamoxifen (Tmx) administration from P1 to P3. NeuN signal identified GC neurons. (**B**) Quantification of proliferating neurons (pH3+; NeuN+) in (A) (n = 5 animals per genotype). (**C**) Quantification of p27 Kip1+ cells in (A) (n = 5 to 9 animals per genotype). (**D-E**) EdU staining of control and *Dab1*^iΔEC^ cerebellar cortices at P10 (D) and P12 (E) after Tmx administration from P1 to P3 and EdU administration at P8. (**F**) Quantification of EdU+ cells per area in each cerebellar cortical layer (n = 7-10 animals per genotype) at P10. (**G**) Distribution of EdU+ cells in (F) per layer. (**H**) Quantification of EdU+ cells per area in each cerebellar cortical layer (n = 5-7 animals per genotype) at P12. (**I**) Distribution of EdU+ cells in (H) per layer. (**J**) anti-Pax6 immunostaining of granule cells in the granule cell layer in 4-week-old mice after Tmx administration from P1-P3. (**K**) Quantification of Pax6+ cells per area in the GCL in (J) (n = 8 animals per genotype). Abbreviations: EGL = external granule layer, ML = molecular layer, GCL = granule cell layer. Scale bars: A, D, E and J = 50 µm. Data are shown as mean ± SEM. \**P*<0.05, \*\**P*<0.01.

To determine whether this increase in proliferation also altered GCP migration or final positioning, we performed EdU birthdating at P8 and analyzed the distribution of labeled cells at P10 and P12. While *Dab1*^iΔEC^ mice displayed a higher density of EdU⁺ cells in the EGL at both timepoints and also in the ML at P12 (Figure 2D-2F and 2H), the relative distribution of cells across the EGL, ML, and GCL was comparable to controls (Figure 2G and 2I), indicating that radial migration was not impaired. Additionally, no ectopic accumulation of Pax6⁺ cells was detected in the ML at 4 weeks of age (Figure S2C and S2D), suggesting that mature GCs reached their correct laminar destinations. Finally, Pax6⁺ cell density in the GCL at 4 weeks was significantly increased in *Dab1*^iΔEC^ mice (Figure 2J and 2K), indicating that the postnatal expansion of the GCP pool results in a persistent increase in granule cell output.

Together, these results demonstrate that endothelial Dab1 restricts GCP proliferation, supporting an instructive role for vascular signaling in postnatal cerebellar neurogenesis.

### A Dab1–Wnt5a signaling axis links blood vessels to granule cell progenitor control

To identify candidate signaling molecules that could mediate the effects of endothelial Dab1 on cerebellar development, we searched for secreted factors downregulated in the absence of Dab1. The non-canonical ligand Wnt5a has been shown to orchestrate cerebellar development from early embryonic stages ^19^ and is also expressed in endothelial cells where it contributes to pro-angiogenic signaling ^20,21^. In brain endothelial cells, Dab1 knockdown led to downregulation of *Wnt5a* mRNA (Figure 3A).

**Fig. 3.**
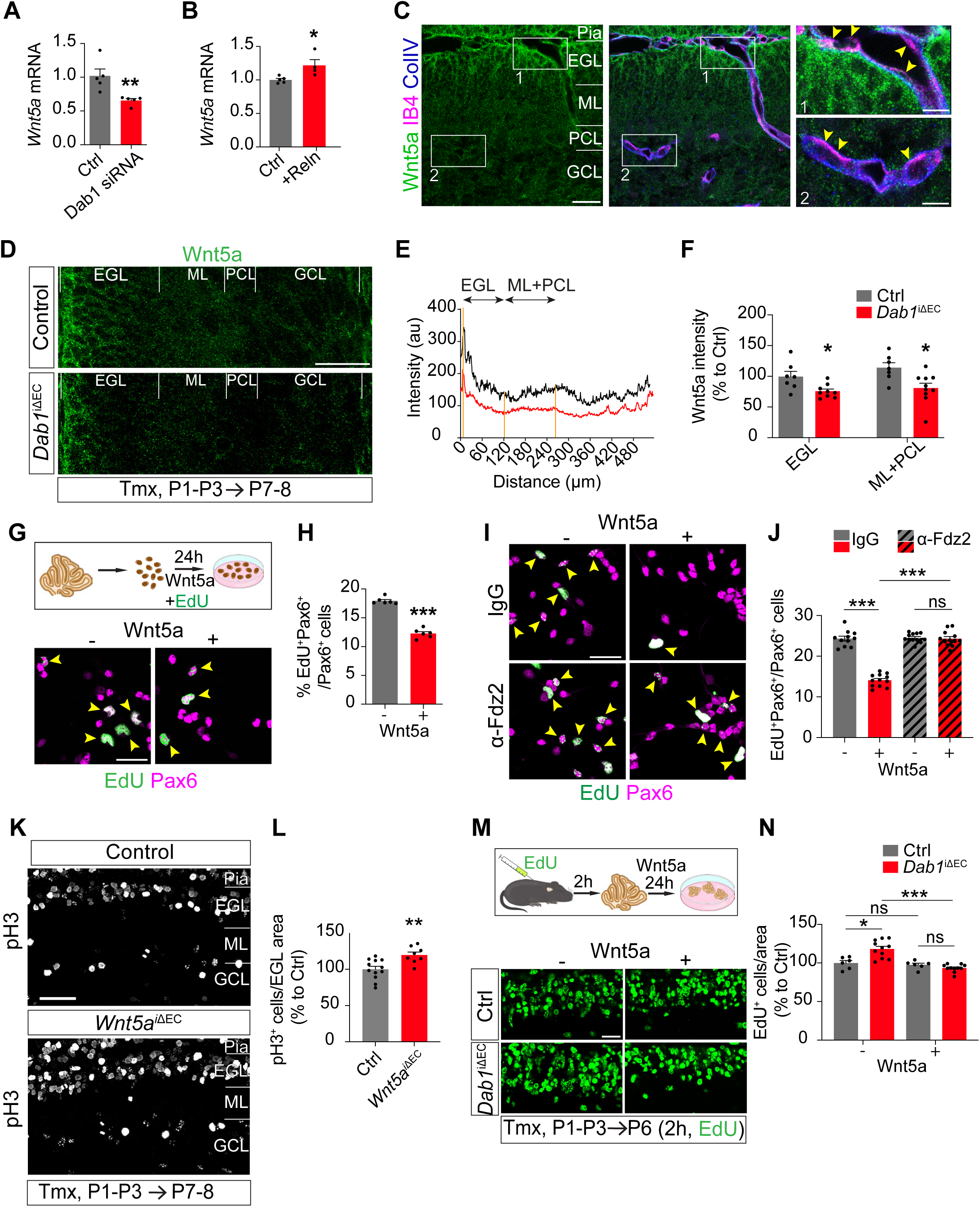
Endothelial Dab1 induces vascular Wnt5a to limit granule cell progenitor proliferation via Fzd2. (**A**) *Wnt5a* mRNA expression is decreased in bEnd.3 cells treated with *Dab1* siRNA compared to siRNA control, assessed with quantitative PCR (n = 5). (**B**) *Wnt5a* mRNA expression is increased in bEnd.3 cells after 24 h stimulation with exogenous Reelin (Reln) protein, assessed by quantitative PCR (n =4-5). (**C**) Wnt5a immunostaining in cerebellar cortex at P8. Rectangles 1 and 2 are magnified on the right. Arrows indicate Wnt5a signal in a pial (1) and a cerebellar cortical vessel below the PCL (2). Blood vessels are detected with IB4 and Collagen IV (ColIV). (**D**) Wnt5a staining of control and *Dab1*^iΔEC^ cerebellar cortices at P8 after tamoxifen (Tmx) administration from P1 to P3. (**E**) Plot profile of Wnt5a signal intensity in (D). (**F**) Quantification of the Wnt5a intensity in the EGL and ML together with the PCL in (D) (n = 7-10 animals per genotype). (**G**) EdU and Pax6 labeling of granule cell progenitors (GCP) primary cell culture stimulated with exogenous Wnt5a. Arrows indicate proliferating GCP (EdU+ Pax6+). (**H**) Quantification of the proportion of proliferating GCP (EdU+ Pax6+) in (G) (n =6). (**I**) EdU and Pax6 labeling of GCP primary cell culture stimulated with exogenous Wnt5a and blockade of the Fzd2 receptor with anti-Fzd2 antibody. Arrows indicate proliferating GCP (EdU+ Pax6+). (**J**) Quantification of the proportion of proliferating GCP (EdU+ Pax6+) in (I) (n =11-12). (**K**) anti-pH3 immunostaining was used to determine the cell cycle stage of granule cell precursors in control and *Wnt5a*^iΔEC^ cerebellar cortices at P8 after Tmx administration from P1 to P3. (**L**) Quantification of proliferating neurons (pH3+) per EGL area in (K) (n = 8-12 animals per genotype). (**M**) EdU staining of P6 control and *Dab1*^iΔEC^ organotypic cerebellar cortices cultured after Tmx administration from P1 to P3 and stimulated with exogenous Wnt5a. (**N**) Quantification of EdU+ cells per EGL area in (M) (n = 6-11 animals per genotype). Abbreviations: EGL = external granule layer, GCL = granule cell layer, ML = molecular layer and PCL = Purkinje cell layer. Scale bars: C, G and I = 30 µm; squares in C =10 µm; D and K = 50 µm; M = 20 µm. Data are shown as mean ± SEM. \**P*<0.05, \*\*\**P*<0.001, ns = non significant.

To validate Wnt5a as a downstream target of Reelin-Dab1 signaling in endothelial cells, we stimulated cultured bEnd.3 cells with exogenous Reelin. *Wnt5a* mRNA expression was significantly upregulated (Figure 3B), and Western blotting confirmed an increase in both major Wnt5a isoforms, long and short (Fig. S3A and S3B). Consistent with these findings, Wnt5a mRNA and protein were detected in cerebellar blood vessels in vivo, particularly in the pial vasculature and in vessels beneath the Purkinje cell layer PCL, as shown by fluorescent in situ hybridization and immunostaining (Figure S3C and Figure 3C). Wnt5a protein displayed a spatial distribution with the highest signal near the pial surface, and this gradient was markedly diminished in *Dab1* mice (Figure 3D). Quantification revealed significantly reduced Wnt5a protein levels in both the EGL and ML-PCL regions in *Dab1*^iΔEC^ animals (Figure 3D and 3E), confirming that endothelial Dab1 is required for Wnt5a expression in the developing cerebellum in vivo.

We next assessed whether vascular Wnt5a directly regulates GCP proliferation. Treatment of primary GCP cultures with recombinant Wnt5a protein resulted in a significant reduction in EdU incorporation among Pax6⁺ GCPs, indicating decreased proliferation (Figure 3F and 3G). This was supported by reduced *Ki67* mRNA expression (Figure S3D). Importantly, TUNEL staining showed no increase in apoptosis following Wnt5a treatment, excluding cell death as a confounding factor (Figure S3E and S3F).

To gain further insight into the mechanism by which endothelial Wnt5a regulates GCP proliferation, we investigated its potential receptor in GCPs. Based on single-nucleus transcriptomic data from the developing mouse cerebellum ^22^, we focused on Fzd2, which is selectively expressed in proliferating GCPs during early postnatal development. Fluorescent in situ hybridization confirmed strong *Fzd2* mRNA expression in GCPs within the EGL and in Purkinje cells (Figure S3G). To test whether Fzd2 mediates the anti-proliferative effects of Wnt5a, we stimulated primary GCP cultures with recombinant Wnt5a while blocking Fzd2 using a specific antibody. Blocking Fzd2 prevented the Wnt5a-induced reduction in EdU incorporation (Figure 3H and 3I), indicating that Wnt5a signals through Fzd2 to limit GCP proliferation.

### Wnt5a secreted by endothelial cells is necessary and sufficient to restrain granule cell progenitor proliferation

To directly test the requirement for vascular Wnt5a as the key downstream effector of endothelial Dab1 signaling in vivo, we generated *Wnt5a*^iΔEC^ mice by crossing *Cdh5-creERT2* mice and *Wnt5a^lox/lox^* mice ^23^. 4OH-Tamoxifen was administered from P1 to P3, and cerebellar tissue was analyzed at P8 and 4 weeks. We then assessed GCP proliferation in *Wnt5a*^iΔEC^ mice, paralleling our analysis in *Dab1*^iΔEC^ animals. Quantification of phospho-Histone H3 (pH3)–positive cells in the EGL revealed a significant increase in mitotic GCPs in *Wnt5a*^iΔEC^ mice compared to littermate controls (Figure 3J and 3K), mirroring the phenotype observed in *Dab1*^iΔEC^ mice (see Figure 2A and 2B). These results demonstrate that vascular Wnt5a is essential for limiting GCP proliferation during early postnatal cerebellar development and acts as a downstream effector of endothelial Dab1 signaling in vivo.

We then asked whether Wnt5a is sufficient to rescue the GCP hyperproliferation observed in *Dab1*^iΔEC^ cerebella. To test this, we cultured organotypic cerebellar slices from *Dab1*^iΔEC^ and control mice, both treated with exogenous Wnt5a protein. As expected, *Dab1*^iΔEC^ slices displayed elevated EdU⁺ GCP density compared to controls. Remarkably, Wnt5a treatment fully restored GCP proliferation to control levels (Figure 3L and 3M), demonstrating that Wnt5a is sufficient to normalize GCP cell cycle dynamics in the absence of endothelial Dab1.

Together, these findings identify Wnt5a as a critical endothelial-derived effector downstream of Reelin-Dab1 signaling and establish its functional role in restraining GCP proliferation during early postnatal cerebellar development.

### Purkinje cell dendritogenesis depends on the Dab1-Wnt5a endothelial signaling axis

At P7-P8, PCs transition from a rounded somata to a polarized morphology ^24,25^ as their dendrites expand into the ML in parallel with the extension of granule cell–derived parallel fibers (PFs). Building on our demonstration of the close alignment between Purkinje cells and blood vessels in the PCL, as well as our identification of Dab1 and Wnt5a in these vessels and Fzd2 expression in Purkinje cells, we examined whether vascular signaling directly contributes to Purkinje cell development. To test this hypothesis, we examined Purkinje cell (PC) morphology in *Dab1*^iΔEC^ mice using calbindin (Cb) immunostaining at P8. Mutant PCs exhibited significantly rounder somata and reduced dendritic complexity, resulting in a shortened nascent molecular layer (ML) (Figure 4A–4C). By 4 weeks of age, PCs in *Dab1* ^iΔEC^ mice showed a significantly reduced distance from the soma to the first dendritic branch point and fewer branch points up to the third-order dendrite compared with controls (Figure 4D–4F), indicating persistent impairments in dendritic maturation.

**Fig. 4.**
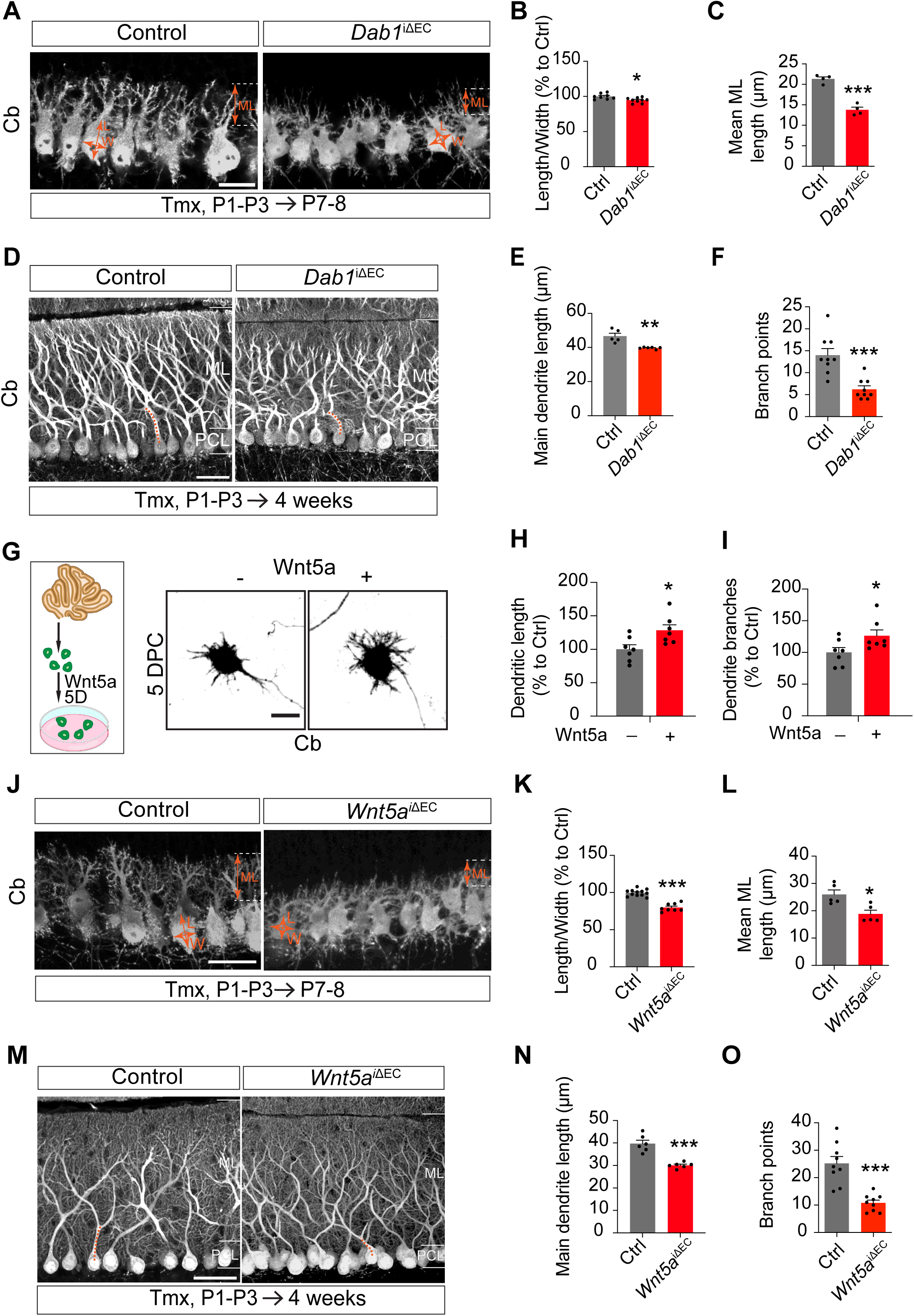
Vascular Wnt5a promotes Purkinje cell dendritic maturation. (**A**) Purkinje cells (PC) stained with anti-Calbindin (Cb) antibody in control and *Dab1*^iΔEC^ cerebellar cortices at P8 after tamoxifen (Tmx) administration from P1 to P3. (**B**) Quantification of length (L)/width (W) PC soma in (A) (n = 8-9 animals per genotype). (**C**) Quantification of the mean molecular layer length in (A) (n = 4 animals per genotype). (**D**) Purkinje cells (PC) stained with an antibody against Calbindin (Cb) in control and *Dab1*^iΔEC^ cerebellar cortices in 4-week-old mice after Tmx administration from P1 to P3. (**E**) Quantification of the distance from the first branch point in the main dendrite to PC soma in D (orange dashed line) (n = 5-6 animals per genotype). (**F**) Quantification of number of branch points measured up to the tertiary dendrite in D (n = 9 neurons, 3 animals per genotype). (**G**) Cb immunostaining of PC primary cell culture stimulated with Wnt5a after 5-days (D) post culture (DPC). (**H**-**I**) Quantifications of the dendritic length (H) and number of dendritic branches (I) in (G) (n = 7-8 neurons). (**J**) Purkinje cells (PC) stained with anti-Calbindin (Cb) antibody in control and *Wnt5a*^iΔEC^ cerebellar cortices at P8 after Tmx administration from P1 to P3. (**K**) Quantification of length (L)/width (W) PC soma in (J) (n = 8-12 animals per genotype). (**L**) Quantification of the mean molecular layer length in (J) (n = 5 animals per genotype). (**M**) Purkinje cells (PC) stained with an antibody against Calbindin (Cb) in control and *Wnt5a*^iΔEC^ cerebellar cortices in 4-week-old mice after Tmx administration from P1 to P3. (**N**) Quantification of the distance from the first branch point in the main dendrite to PC soma in M (orange dashed line) (n = 6 animals per genotype). (**O**) Quantification of number of branch points measured up to the tertiary dendrite in M (n = 9 neurons, 3 animals per genotype). Abbreviations: ML = molecular layer, PCL = Purkinje cell layer. Scale bars: A = 20 µm; D and M = 50 µm; G and J = 30 µm. Data are shown as mean ± SEM. \**P*<0.05, \*\**P*<0.01, \*\*\**P*<0.001.

We next asked whether the same Dab1–Wnt5a vascular axis operates during Purkinje cell dendritogenesis. To assess whether Wnt5a directly promotes PC dendritic development, we cultured primary PCs from C57BL/6 mice and treated them with recombinant Wnt5a. Wnt5a-treated PCs exhibited increased expansion in dendritic complexity, with significantly more branches and increased arbor total length (Figure 4G-4I). Importantly, we were able to recapitulate in vivo the PCs dendritic arbor defects in endothelial Wnt5a mutants. At P8, calbindin-labeled PCs in *Wnt5a*^iΔEC^ mice displayed significantly rounder somata and less elaborated dendrites compared to controls (Figure 4J–4L) and at 4 weeks of age, this morphological impairment persisted (Figure 4M-4O), establishing that endothelial Wnt5a is both sufficient and necessary for proper Purkinje cell dendritic maturation during postnatal development.

### Loss of vascular signaling weakens Purkinje-cell firing and plasticity

To investigate whether morphological abnormalities observed in *Dab1*^iΔEC^ Purkinje cells were accompanied by alterations in their physiological function, we examined spontaneous and evoked firing properties at 4 weeks of age, a stage when PC electrophysiological features are considered mature ^26^, and directly comparable to our morphological data.

PCs fire spontaneous simple spikes driven by intrinsic ion channels ^6,27^. Using a cell-attached recording protocol, we measured spontaneous tonic activity and found that *Dab1*^iΔEC^ PCs displayed a significantly reduced firing frequency compared to control PCs (Figure 5A–5D). To determine whether intrinsic excitability was affected, we performed current-clamp recordings with input-output current injection. We found no significant difference between *Dab1*^iΔEC^ and control mice in either firing threshold or spike response to current injection (Figure 5E–5G), suggesting that the observed firing reduction is not due to intrinsic membrane property changes.

**Fig. 5.**
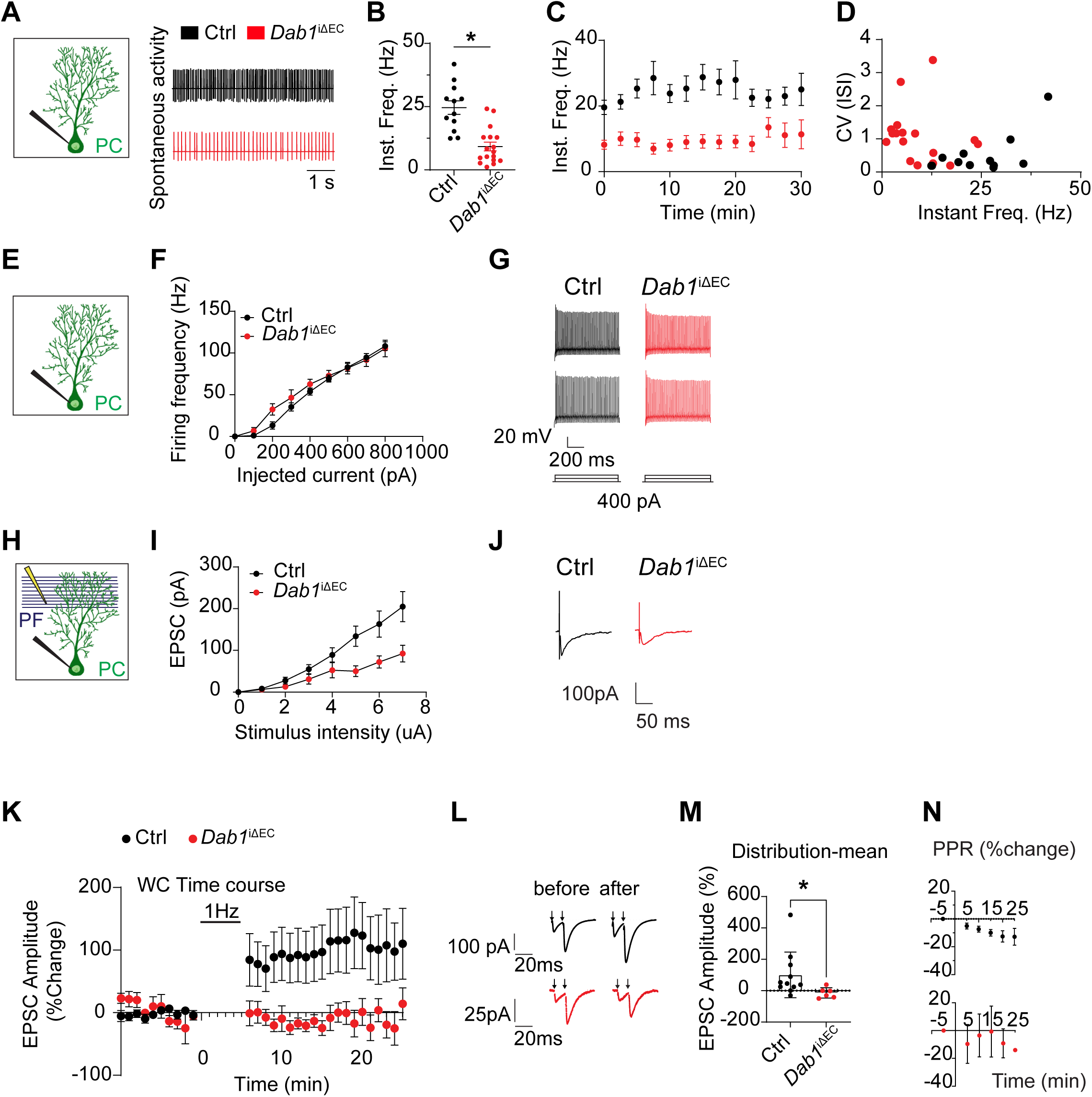
Loss of endothelial Dab1 impairs Purkinje cell firing and parallel fiber synaptic plasticity. (**A-D**) PC of *Dab1*^iΔEC^ and control mice were patched at 3-4 weeks of age after tamoxifen (Tmx) administration from P1 to P3 and spontaneous tonic firing was recorded (see schematic representation) (n = 12-17 PC per genotype). Examples of traces are shown in (A). Instantaneous frequency was analyzed in (B-D). (**E-G**) PC of *Dab1*^iΔEC^ and control mice were patched at 3-4 weeks of age after Tmx administration from P1 to P3 and intrinsic excitability was assessed (see schematic representation in E) (n = 10-15 PC per genotype). Current-frequency plot shown in (F) and example of single traces with action potentials evoked in a single step pulse in (G). (**H-J**) PC from (E-G) and parallel fibers (PF)-PC synaptic output was recorded (see schematic representation). Summary plot of the average amplitude of excitatory postsynaptic current (EPSC) to increasing intensity of stimuli in (I). Example of single EPSC in (J). (**K-N**) PC of *Dab1*^iΔEC^ and control mice were patched at 3-4 weeks of age after Tmx administration from P1 to P3 and synaptic plasticity at PF-PC synapses was recorded (see schematic representation in H) (n = 6-11 PC per genotype). Induction of long-term potentiation (LTP) at PF-PC synapse after 1 Hz for 5 min stimulation in (K). Examples of single traces in (L). Average of EPSC amplitude after 20 min from the induction (% change to control) in (M). Pair-pulse ratio (PPR) at the PF-PC synapse after LTP induction in (N). Abbreviations: PC = Purkinje cell, PF = parallel fibers. Data are shown as mean ± SEM. \**P*<0.05.

We next assessed whether synaptic input from granule cells was impaired. Granule cell axons, known as parallel fibers (PF), form glutamatergic synapses onto PC dendrites ^3^. To examine PF–PC connectivity, we stimulated PFs and recorded the evoked EPSCs in PCs. *Dab1*^iΔEC^ PCs exhibited a significant reduction in PF-evoked EPSC amplitude (Figure 5H-5J), suggesting compromised synaptic transmission. To determine whether synaptic plasticity was also affected, we induced long-term potentiation (LTP) at the PF–PC synapse. In contrast to control PCs, which exhibited robust synaptic potentiation, *Dab1*^iΔEC^ PCs showed markedly impaired LTP induction (Figure 5K–5N). These results indicate that endothelial Dab1 signaling is required for both basal synaptic transmission and activity-dependent synaptic strengthening at PF–PC synapses.

### Excitatory synapse assembly onto Purkinje cells requires vascular cues

Given the impaired PF–PC transmission and plasticity observed in *Dab1*^iΔEC^, we next asked whether these functional deficits could be explained by alterations in PF axon organization and PF–PC synaptic architecture.

To visualize PF organization in the cerebellar cortex, we applied DiI crystals to the cerebellar surface and incubated the tissue for 6 days to allow anterograde diffusion through granule cell axons. In control animals, PFs displayed highly parallel and coherently aligned trajectories within the molecular layer (ML), as expected. In contrast, *Dab1*^iΔEC^ mice exhibited disorganized PF architecture, with misaligned axons and reduced directional coherence. Quantification using the coherency coefficient, a measure of fiber orientation consistency ^28^, revealed a significant decrease in PF alignment in *Dab1*^iΔEC^ mice (Figure 6A and 6B), indicating that endothelial Dab1 signaling is required for proper spatial organization of PF projections.

**Fig. 6.**
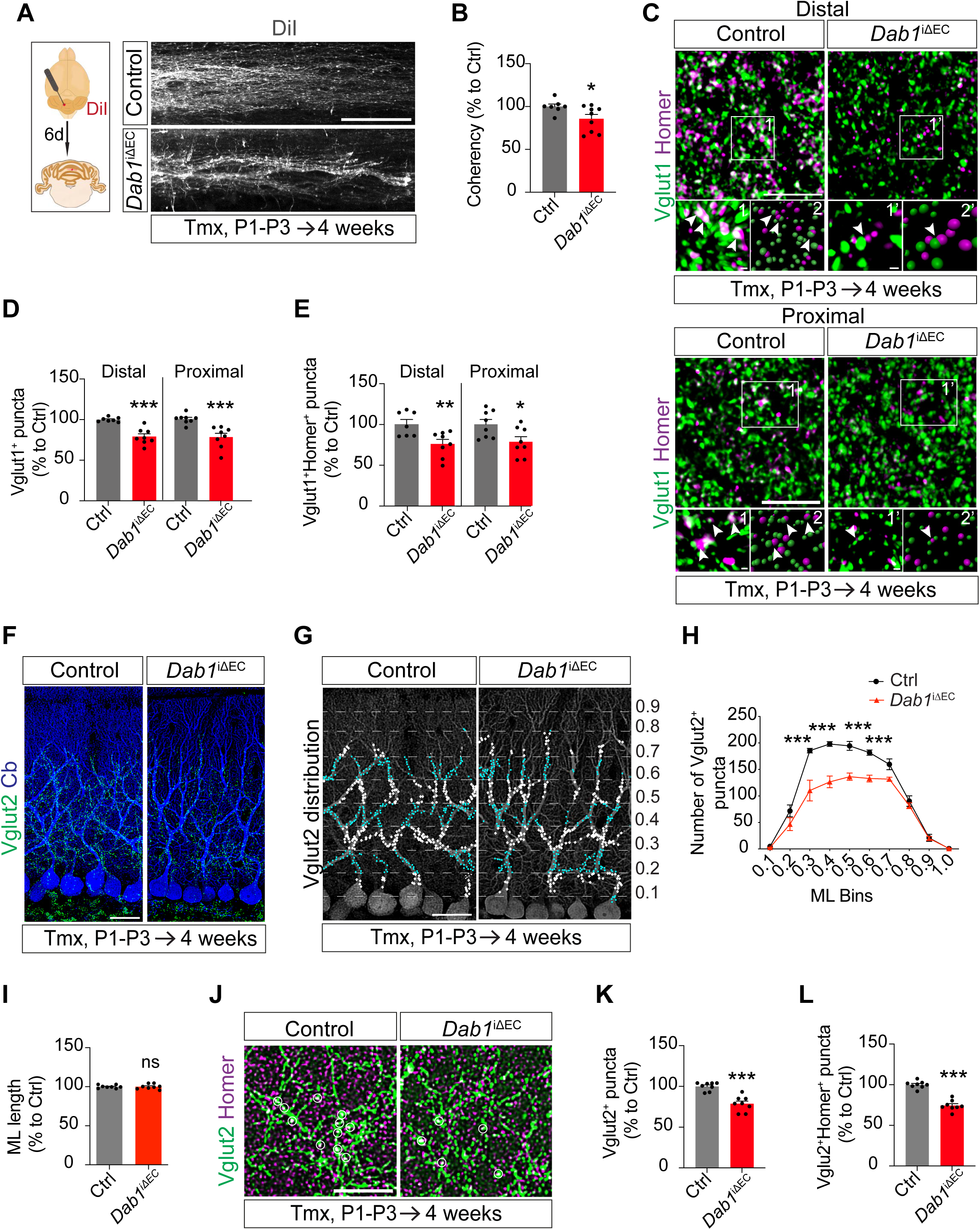
Endothelial Dab1 signaling is required for excitatory synapse assembly onto Purkinje cells. (**A**) DiI staining of granule cell parallel fibers in the molecular layer (ML) in 4-week-old mice after tamoxifen (Tmx) administration from P1-P3. (**B**) Quantification of parallel fiber coherency in (A) (n = 8 animals per genotype). (**C**) Parallel fiber terminals stained with anti-Vglut1 antibody and post-synaptic element in Purkinje cells (PC) detected with anti-Homer antibody in distal and proximal part of PC dendritic arbors in *Dab1*^iΔEC^ and control cerebellar cortices in 4-week-old mice after Tmx administration from P1 to P3. Higher magnifications (1 and 1’) and corresponding IMARIS reconstructions (2 and 2’) are enlarged below. Arrows indicate co-localization of Vglut1^+^Homer^+^ puncta. (**D**) Quantification of Vglut1+ and Vglut1+ Homer+ puncta in (C distal and proximal) (7-8 animals per genotype). (**F-H**) Climbing fiber terminals immunolabelled with anti-Vglut2 antibody and post-synaptic element in PC (identified with calbindin (Cb)) detected with anti-Homer antibody in *Dab1*^iΔEC^ and control cerebellar cortices in 4-week-old mice after Tmx administration from P1 to P3 (n = 8 animals per genotype). Distribution of Vglut2+ puncta from (F) throughout the binarized PC dendritic arbor (G). Quantification of the distribution of Vglut2+ puncta from each bin division in the ML (H). (**I**) Quantification of ML distance from Purkinje cell body to pial surface (n = 8 animals per genotype). (**J**) Immunostaining with anti-Vglut2 and anti-Homer. Circles indicate the co-localization of Vglut2+ Homer+ puncta. (**K-L**) Quantification of Vglut2^+^ (K) and Vglut2^+^Homer^+^ puncta (L) in (J). Scale bars: A = 50 µm; C and J = 5 µm; magnified regions in C = 0.5 µm; F-G = 10 µm. Data are shown as mean ± SEM. \**P*<0.05, \*\**P*<0.01, \*\*\**P*<0.001, ns = non significant.

We next assessed whether this disruption in PF organization was accompanied by defects in PF–PC synapse formation. Immunostaining for the vesicular glutamate transporter Vglut1, which selectively labels PF presynaptic terminals in the ML, together with the postsynaptic scaffold protein Homer, was used to quantify mature excitatory synapses. We separately quantified synaptic markers in both distal and proximal regions of the PC dendritic arbor, given that PFs primarily target distal dendrites, whereas proximal dendrites receive a mix of PF and climbing fiber (CF) inputs ^3,29^. In both distal and proximal dendrites, *Dab1*^iΔEC^ mice displayed a significant reduction in Vglut1^+^ puncta, indicating fewer PF presynaptic terminals (Figure 6C and 6D). Moreover, Vglut1^+^Homer^+^ puncta — representing active glutamatergic PF–PC synapses — were also markedly reduced in both regions, with the effect more pronounced in distal dendrites (Figure 6C and 6D), consistent with the known anatomical distribution of PF inputs.

During postnatal cerebellar development, PFs and CFs compete for synaptic territory on the developing PC dendritic arbor ^29^. In several ataxia models and synaptic plasticity mutants, loss of PF–PC synapses is associated with increased CF innervation, suggesting compensatory remodeling ^30^. To determine whether a similar reorganization occurred in our vascular mutants, we examined CF synaptic architecture in *Dab1*^iΔEC^ mice at 4 weeks of age.

Immunostaining for vesicular glutamate transporter 2 (Vglut2), a marker specific to CF presynaptic terminals, revealed no gross alterations in the overall distribution or territorial extent of CF innervation along the PC dendritic tree when comparing *Dab1*^iΔEC^ mice and littermate controls (Figure 6F-6H). Consistent with this, ML thickness was unchanged between genotypes, indicating that global dendritic expansion and laminar organization were preserved (Fig. 6I). However, quantitative analysis revealed a significant reduction in the density of Vglut2⁺ puncta in *Dab1*^iΔEC^ mice, indicating fewer CF terminals contacting the PC dendrites (Fig. 6G, 6H, 6J and 6K). Importantly, Vglut2^+^ Homer^+^ colocalized puncta were also significantly reduced, demonstrating a loss of functional CF–PC synapses (Figure 6J and 6L).

Together, these findings indicate that endothelial Dab1 signaling is required not only for PF–PC synaptic development, but also for the proper establishment or maintenance of CF–PC connectivity. The concomitant reduction of both major excitatory inputs onto PCs provides a structural basis for the impaired firing properties and synaptic plasticity observed in *Dab1*^iΔEC^ mice. These results underscore a central role for vascular-derived cues in coordinating excitatory synapse assembly and circuit integration across multiple afferent systems in the cerebellar cortex.

## Discussion

In this study, we identify a vascular-to-neuronal signaling axis that orchestrates postnatal cerebellar development through direct communication between endothelial cells and neurons. We show that the neuronal guidance molecule Reelin, expressed predominantly by granule cells in the cerebellum, activates Dab1 signaling in endothelial cells, in turn inducing vascular expression and secretion of the non-canonical Wnt ligand Wnt5a. Endothelial-derived Wnt5a restrains granule cell progenitor (GCP) proliferation and promotes Purkinje cell (PC) dendritic maturation. Disruption of this pathway during early postnatal development leads to persistent structural and functional defects in cerebellar circuitry, including abnormal parallel fiber organization, reduced excitatory synapse formation, and impaired synaptic plasticity in young adult animals. Together, these findings establish blood vessels as active, instructive regulators of cerebellar circuit assembly.

The cerebellum is particularly well suited to reveal instructive roles of the vasculature, as much of its neuronal development occurs postnatally, in temporal synchrony with angiogenic expansion. While previous work in the cerebellum has primarily linked vascular function to oxygen delivery and metabolic control of neurogenesis ^12^, our data demonstrate that blood vessels also provide molecular signals that actively instruct neuronal behavior. We found that cerebellar angiogenesis follows a highly structured spatial and temporal pattern, with vascular branching closely aligned to emerging neuronal layers and circuits. This organization is disrupted in endothelial Dab1 mutants, indicating that Reelin–Dab1 signaling contributes to proper vascular patterning and suggesting that angiogenesis itself may be an integral component of cerebellar circuit formation.

Reelin is a central regulator of layered CNS structures, and its loss results in profound cerebellar malformations characterized by disrupted lamination and hypoplasia ^14–16,31^. Beyond its neuronal functions, we have previously shown that Reelin also exerts pro-angiogenic effects in the CNS. In the cerebral cortex, endothelial Dab1 signaling regulates laminin deposition and radial glial anchorage at the pial surface, thereby controlling neuronal migration and blood–brain barrier formation ^10^. Importantly, the vascular role of Dab1 uncovered here is mechanistically and temporally distinct from its previously described functions in cortical development. Despite increased granule cell progenitor proliferation, granule cell migration and laminar positioning were largely preserved following endothelial Dab1 deletion, in contrast to the cerebral cortex where neuronal migration is disrupted. In the cerebellum, proliferation within the external granule layer is temporally separated from inward migration, and this migratory step remains intact in endothelial Dab1 mutants, Instead, vascular Dab1 in the cerebellum acts as a signaling hub that drives secretion of the non-canonical Wnt ligand Wnt5a, which directly regulates GCP proliferation, PC dendritogenesis, and excitatory synapse assembly. This shift highlights a mechanistically distinct role for vascular signaling in the cerebellum, where endothelial cells act as sources of instructive cues that regulate neuronal growth and circuit assembly. These findings indicate that endothelial Dab1 signaling is context-dependent and that the vasculature deploys distinct molecular outputs to regulate successive stages of neural development across brain regions.

Mechanistically, our work establishes Wnt5a as a key endothelial-derived effector downstream of Reelin–Dab1 signaling in the cerebellum. Wnt5a has well-established roles in neuronal development, including axon guidance, dendritic growth, and synapse formation in multiple brain regions ^32–35^. During embryonic cerebellar development, loss of Wnt5a or pan-Wnt secretion leads to hypoplasia and impaired generation of both glutamatergic and GABAergic neurons ^19,36^. However, conditional deletion of Wnt5a from Nestin-positive neuronal progenitors affects PC maturation without altering GCP proliferation or synaptic connectivity ^19^, suggesting that neuronal Wnt5a is dispensable for key postnatal developmental processes. By contrast, our results demonstrate that the vascular source of Wnt5a is essential for postnatal cerebellar development. Endothelial Wnt5a deletion phenocopies endothelial Dab1 loss, while exogenous Wnt5a restores normal GCP proliferation in Dab1-deficient cerebellar slices, establishing both necessity and sufficiency. These findings identify the vasculature as a previously unappreciated source of Wnt5a that regulates neurogenesis and neuronal maturation during a critical developmental window.

Spatially, our data reveal that endothelial-derived Wnt5a is not uniformly distributed throughout the cerebellar cortex, but instead forms a gradient emanating from pial and sub-PCL vessels toward the external granule layer. This spatial organization suggests a locally acting signaling mechanism in which Reelin, secreted by postmitotic granule cells adjacent to the pial surface, activates Dab1 in neighboring meningeal and penetrating vessels to induce Wnt5a release. In this model, vascular Wnt5a would act in a paracrine manner to constrain proliferation of granule cell progenitors positioned immediately beneath the pial vasculature, thereby coupling the local cellular composition of the EGL to vascular-derived signals. Such spatially restricted signaling provides a mechanism by which progenitor pool size can be precisely tuned in accordance with laminar maturation, rather than being governed by solely global cues. Consistent with this idea, components of the Wnt secretion machinery, including Wntless, have been reported in the meninges during cerebellar development ^37^, supporting the concept that meningeal and vascular compartments serve as active signaling interfaces rather than passive barriers. This spatial mode of regulation further suggests that Wnt5a signaling in the cerebellum is highly context dependent, with distinct sources exerting non-redundant functions across developmental time. Previous work has shown that Wnt5a is expressed by the choroid plexus epithelium and secreted into the cerebrospinal fluid during embryogenesis, where its loss results in reduced cerebellar size without overt effects on proliferation or apoptosis ^38^. Together with our findings, this supports a model in which early, long-range Wnt5a signals contribute to gross cerebellar morphogenesis, whereas later, vascular-derived Wnt5a acts locally to regulate postnatal progenitor dynamics and neuronal maturation. The ability of the same morphogen to exert distinct developmental effects depending on its cellular source, spatial range, and receptor context underscores the importance of tightly controlled ligand delivery in the developing brain.

Importantly, our identification of Frizzled-2 as a receptor mediating Wnt5a-dependent control of GCP proliferation provides a mechanistic link between vascular signaling and progenitor cell-cycle regulation. Fzd2 expression is enriched in proliferative GCPs during early postnatal development and declines as the EGL dissipates, aligning temporally with the window during which vascular Wnt5a exerts its effects. This suggests that endothelial Wnt5a signaling is selectively interpreted by progenitor populations that are competent to respond, thereby preventing inappropriate signaling in more mature neuronal populations. Such receptor-restricted responsiveness may represent a general strategy by which vascular-derived morphogens achieve cell-type specificity within densely packed neural tissues.

Beyond its effects on granule cell progenitors, our findings reveal a direct role for vascular signaling in Purkinje cell dendritogenesis and synaptic maturation. Purkinje cells are traditionally thought to undergo largely cell-autonomous dendritic development during the first postnatal week, with later refinement driven by synaptic interactions with granule cells and climbing fibers ^25,39,40^. Our findings identify a vascular signaling mechanism in which endothelial cells translate neuronal cues into morphogenetic outputs that coordinate postnatal cerebellar circuit assembly. The close alignment of Purkinje cell somata with a vascular plexus beneath the PCL positions these neurons to directly receive vascular cues, and the expression of the Wnt receptor Fzd2 in Purkinje cells supports their competence to respond to endothelial-derived Wnt5a signaling. Thus, blood vessels emerge as important regulators of Purkinje cell intrinsic maturation programs.

As development proceeds, the consequences of early vascular signaling deficits propagate across multiple levels of cerebellar circuit organization. Loss of endothelial Dab1 or Wnt5a not only compromises Purkinje cell dendritic architecture, but also disrupts the orderly alignment of parallel fibers, reduces excitatory synapse formation from both parallel and climbing fibers, and weakens synaptic transmission and plasticity. These defects span distinct stages of circuit assembly, from axonal organization and synapse formation to activity-dependent refinement, indicating that vascular signaling acts upstream of multiple developmental processes rather than affecting a single node in the circuit. The concomitant impairment of both major excitatory afferent systems onto Purkinje cells is particularly informative. Parallel fibers and climbing fibers differ in their developmental timing, anatomical targeting, and modes of synaptic plasticity, yet both are reduced in Dab1 vascular mutants. This argues against a simple secondary effect driven by increased granule cell number or delayed neuronal maturation and instead suggests that vascular-derived cues help establish a permissive structural and molecular framework that supports convergent synaptic integration onto Purkinje cells.

Importantly, neuronal deletion of Wnt5a does not recapitulate the synaptic and circuit-level defects observed here ^19^, underscoring that vascular-derived Wnt5a fulfills a unique and non-redundant role in postnatal cerebellar wiring. This distinction highlights how the source of a developmental signal, rather than the signal itself, can dictate its functional outcome within a circuit. Together, our findings support a model in which endothelial cues shape the structural and functional landscape of the cerebellar cortex, thereby enabling the proper emergence of synaptic connectivity and plasticity in the mature circuit.

### Broader implications and future directions

Together, our findings support a model in which endothelial cells act as developmental organizers that integrate neuronal-derived signals and translate them into morphogenetic outputs regulating progenitor behavior, neuronal maturation, and synaptic integration. This work uncovers a multistep, cell-type-dependent neurovascular communication axis, in which neuronal cues instruct endothelial signaling programs that in turn shape postnatal cerebellar circuit assembly.

More broadly, our study suggests that angiogenesis and neural circuit assembly are mechanistically intertwined processes, coordinated through reciprocal signaling interactions. Given that Purkinje cell dysfunction underlies a wide range of neurological disorders, including inherited ataxias and neurodevelopmental conditions, subtle disruptions in vascular signaling during development may contribute to disease susceptibility by perturbing circuit assembly rather than causing overt malformations. Understanding how vascular-derived signals shape neuronal development may therefore open new avenues for therapeutic intervention, leveraging the vasculature as an accessible and modulatable interface to influence neural circuit formation and function.

## Material and methods

### Mice

Animals were bred and maintained in our animal facility in individually ventilated cages under specific germ-free conditions, in 12 hours day/night light cycles, temperature 22 °C, humidity 55%, Aspen bedding and with food and water ab libitum. Experiments were carried out under the approval of the Regierungspräsidium of Darmstadt and the Veterinäramt of Frankfurt am Main.

Endothelial specific Dab1 knockout mice (*Dab1*^iΔEC^) ^10^ and endothelial specific Wnt5a knockout mice (*Wnt5a*^iΔEC^) were generated both by crossing *Cdh5-creERT2* mice (kindly provided by R. Adams) and *Dab1^lox/lox^* (kindly provided by U. Müller) or *Wnt5a^lox/lox^* ^23^ (Jackson Laboratory Strain #026626) were used in the present study. Cre activity was induced by intraperitoneal injection of 50 µl (1 mg/ml) of 4-hydroxytamoxifen (4OHT, Tmx) each day for 3 days (P1 to P3). Tmx injectable solution was prepared according to ^10^; genotyping and Tmx treatment efficiency was demonstrated in the same publication. *Wnt5a*^iΔEC^ genotyping was performed following the indications of distributor and donating investigators. Cre-negative lox/lox littermates were used as controls, which were processed and analyzed always following the same conditions. For wild-type expression analyses, GCP and PC culture, C57BL/6J mice were used. Both mice genders were used for all the experiments.

### bEnd.3 culture and stimulation

bEnd.3 cells were cultivated in culture medium (DMEM (Dulbecco’s modified Eagle’s medium), 10% fetal bovine serum (FBS), 0.1mg/ml streptomycin, 100U/ml penicillin) at 37 °C, 5% CO_2_. For the stimulation, bEND.3 cells were starved in DMEM for 2 h. Stimulation was performed by adding recombinant Reelin protein (100 ng/ml, R&D, 3820-MR-025) for 24 or 48 h.

For siRNA-mediated Dab1 knockdown, 50–70% confluent bEnd.3 cells were prewashed once with OptiMEM and subsequently treated with siRNA probes at a final concentration of 67 nM mixed with Lipofectamine RNAiMax transfection reagent (13778939, Invitrogen) in OptiMEM for 4 hours at 37 °C. siRNA probes used: *Dab1*:SASI_Mm01_00035228 (Merck) Firefly luciferase (customized as control): CGUACGCGGAAUACUUCGA[dT][dT] (sense) and UCGAAGUAUUCCGCGUACG (antisense). Wnt5a downregulation was assessed by RT-qPCR 48 hours after transfection.

### GCPs culture and treatment

Cerebellar GCPs from P5 C57BL/6 mice were prepared as described previously ^41^. Briefly, cerebellar tissue was dissected and enzymatically dissociated in 20 U/ml of activated papain solution plus 200U/ml of DNase in EBSS (Earle’s Balanced Salt Solution) (Dissociation kit, Worthington) at 37 °C for 15 min. The tissue was triturated and centrifuged. Cell pellets were suspended in EBSS containing 1mg/ml Albumin-Ovomucoid proteinase inhibitor (Dissociation kit, Worthington) and 200U/ml of DNase. The cell suspension was transferred to 10 mg/ml Albumin-Ovomucoid proteinase inhibitor solution, followed by centrifugation to remove debris. To purify the GCPs, cell pellets were suspended in a plating medium NBM (Neurobasal-A medium), 250 μM KCl, 2mM Glutamax, 0.1 mg/ml streptomycin, 100 U/ml penicillin) containing 10% FBS and filtered through a 70 μm strainer to obtain a single-cell suspension. Cells were cultivated on a 60 mm petri-dish coated with 100 μg/ml of poly-D-Lysine in an incubator at 37 °C for 20 min. Only non-adherent cells were collected. After centrifugation, cells were cultivated in culture media (plating media, 1% B27 supplement) on the glass coverslips coated with 500 μg/ml of poly-D-Lysine in an incubator at 37 °C with 5% CO_2_ for 22 h.

Cultured GCP were then stimulated with recombinant Wnt5a protein (100 ng/ml, R&D, 645-WN) and incubated for 24 hours at 37 °C in the presence of 5-ethynyl-2’-deoxyuridine (EdU, Invitrogen, C10337).

To block Frizzled-2 (Fzd2) interaction domain to Wnt5a, cells were treated with culture media containing anti-Fzd2 antibody (rat 20 μg/ml, R&D, MAB1307) or rat IgG isotype control (20 μg/ml, Invitrogen) for 2 h. Afterwards, EdU and Wnt5a or BSA solution were added to the cell culture and incubated for 24 h at 37 °C.

### PC culture and treatment

Primary PC culture was obtained from mouse embryos at embryonic day 17-18. Pregnant females were sacrificed via cervical dislocation and embryos were collected in ice-cold HBSS solution. Cerebellum were removed from embryos and dissociated in 20 U/ml of activated papain solution plus 200 U/ml of DNase in EBSS (Earle’s Balanced Salt Solution) (Dissociation kit, Worthington) at RT for 10 min. Digestion was stopped by adding a papain stop solution (HBSS, 18.25% FBS). After centrifugation, cerebella were collected and triturated in the DNase solution (HBSS, 10 U/ml DNase (Worthington)). The cell suspension was transferred through a 40 μm strainer to obtain a single-cell solution, followed by centrifugation. Cell pellets were suspended in the seeding medium (DMEM, 100 μM putrescine, 30 nM Na_2_CO_3_, 0.1 mg/ml streptomycin,100 U/ml penicillin) with 10% FBS. Cells suspensions were plated on the coverslips coated with poly-D-Lysine and recovered for 3 h at 37 °C, and changed to culture medium (seeding medium, 4 μM progesterone, 0.5 ng/ml triiodothyronine, 25 ng/ml insulin-like growth factor I, 200 μg/ml transferrin, 20 μg/ml insulin). Cells were cultivated for 24 h at 37 °C and replaced by fresh culture medium with anti-Fdz2 antibody (rat 20 μg/ml, R&D, MAB1307) or rat IgG isotype control (20 μg/ml, Invitrogen). 2 hours later, Wnt5a (100 ng/ml, R&D, 645-WN) was added to the cell culture and incubated for 3 or 5 days at 37 °C.

For immunostaining, cells were fixed in 4% PFA for 20 min at RT. Permeabilization was performed in 0.3% Triton X-100 in PBS for 30 min at RT, followed by blocking in 10 % NDS in PBS for 30 min at RT. Primary antibody anti-Calbindin (1:200, Merck, PC253L) was diluted in 2% NDS in PBS and cells were incubated at 4 °C overnight. Incubation in the secondary antibody donkey polyclonal anti-rabbit Alexa Fluor-555 (Invitrogen A31572,1:200) and DAPI was during 1 h at RT.

### Organotypic cerebellar culture

Organotypic cerebellar slice cultures (OTC) were prepared from pups at P6 from *Dab1*^iΔEC^ and control mice. All OTC from the same litter were dissected and processed in the same experimental session on the same day. Isolated cerebella were embedded in 3.5% low-melting agarose. 300 μm sagittal slices were sectioned using a vibratome in cold preparation medium (MEM (modified Eagle’s medium), 25mM HEPES (4-(2-hydroxyethyl)-1-piperazineethanesulfonic acid), 2 mM Glutamax, 0.45% D-Glucose, 0.1 mg/ml streptomycin, 100 U/ml penicillin) at pH 7.4. 6 slices containing cerebellar vermis per mouse were collected and equally divided into two 35 mm petri-dish containing culture medium (42% MEM, 25% basal medium eagle (BME), 25% normal horse serum, 25 Mm HEPES, 0.65% D-glucose, 0.15% NaHCO_3_, 0.1 mg/ml streptomycin, 100 U/ml penicillin, 2 mM Glutamax, 1% N-2 supplement, 1% B-27 supplement), which were cultivated in an incubator at 35 °C. After 1 h of recovery, OTC were transferred on a cell-culture insert (Millipore) and cultivated in the culture media containing 100 ng/ml of Wnt5a at 35 °C for 24 h. Then OTC were fixed in 4% PFA overnight at 4 °C with gentle rotation. After extensive washing in PBS, OCT were cryoprotected and sliced in a cryostat as indicated below.

### DiI Labeling

4-week-old mice were perfused with 4% PFA as described above. Crystals from lipophilic dye 1,1’-Dioctadecyl-3,3,3’,3’-Tetramethylindocarbocyanine Perchlorate (DiI) (D282, Invitrogen) were placed on the cerebellar folia and incubated for 6 days at 4 °C. Cerebella were afterwards post-fixed in 4% PFA at RT for 2 h and sectioned with the vibratome in coronal orientation at 80 µm. Slices were incubated in DAPI-PBS solution and mounted.

### EdU labeling

EdU was reconstituted in 0.9% saline (Invitrogen, C10337). Mice were injected with EdU at 50mg/kg into the P6 or P8 *Dab1*^iΔEC^ and control mice, which were sacrificed 2 h later and used for OTC culture, and at P10 or P12 for GCP migration experiments. Cultured primary GCP were incubated with EdU (2.5 μg/ml) during 24 h together with the stimulation treatments. Then they were fixed in 4% PFA for 20 min at RT. OCT sections, vibratome sections and cultured GCP were permeabilized in 0.5% Triton X-100 in PBS for 20 min at RT and incubated with Click-IT EdU Imaging Kit (Invitrogen, C10337) for 30 min at RT. OCT sections were later incubated with DAPI during 10 min and mounted. Vibratome sections and cultured GCP were immunolabeled as described in the following section.

### Immunostaining and FISH

Animals were deeply anesthetized via intraperitoneal injection of ketamine (100 mg/kg weight, Ketavet) and xylazine (10 mg/kg weight, Rompun) and perfused intracardially with 4% paraformaldehyde (PFA) in phosphate buffered saline (PBS) or 10% TCA. Cerebellum were dissected out and postfixed in 4% PFA or 10% TCA 2 h at room temperature (RT). Cerebella were sectioned sagittal at 80 µm in a vibratome. Alternatively, when cryostat was used for sectioning, cerebellums were cryoprotected by consecutive immersions in 15% and 30% sucrose in PBS at 4 °C and then embedded in Tissue-Tek O.C.T. compound and frozen on dry ice. 16 µm were acquired.

For immunostaining, sections were incubated in blocking-permeabilizing solution (10% NDS, 0.3% Triton X-100 in PBS) during 1 h at RT. Primary antibodies: anti-Reelin (R&D, Af3820), anti-Pax6 (BioLegend, 901301 or Santa Cruz, Sc-81649), anti-Podocalyxin (R&D, AF1556), anti-phospho-Histone H3 (Merck, 06-570), anti-NeuN (Merck, MAB377 or ABN78), anti-p27 Kip1 (Cell Signaling, 83630), anti-Ki67 (Abcam, ab15580), anti-Wnt5a (LS Bio, LS-C124811), anti-Collagen 4 (Bio-Rad, 2150-1470), anti-Glut1 (Millipore, 07-1401), anti-Calbindin D-28k (Swant, 300, Merck, PC253L, Nitobe Medical, AB-2571569), anti-Vglut1(Merck, AB5905), anti-Vglut2 (Frontier Institute, AB_2571621), anti-Homer 1/2/3 (Synaptic Systems, 160 103), anti-Fzd2 (LSBio, LS-B3963) were diluted at 1:200 in 2% NDS, 0.1% Triton X-100 in PBS and incubated during 48 h at 4 °C. After consecutive washes in 0.1% Tween 20 in PBS, secondary antibodies: goat anti-Guinea pig A488 (Life Technologie, A11073), goat anti-rabbit A568 (Invitrogen, A32732), goat anti-mouse A647 (Invitrogen, A32728), donkey anti-mouse A488 (Invitrogen, A21202), A568 (Invitrogen, A10037), A647 (Invitrogen, A31571), donkey anti-rabbit A488 (Invitrogen, A21206), A555 (Invitrogen, A31572), A647 (Invitrogen, A31573), donkey anti-goat A488 (Invitrogen, A11055), A568 (Invitrogen, A11057), A647 (Invitrogen, A21447), donkey anti-rat A488 (Invitrogen, A21208), were diluted at 1:200 in the same solution and incubated overnight at 4 °C. To visualize cell nuclei and blood vessels, DAPI (1:1000) and IB4 (1:200, Isolectin GS-IB4 from *G. simplicifolia* Alexa Fluor-568, Thermo Fisher, 21412) was added in the secondary antibody incubation solution. Afterwards sections were washed in 0.1% Tween 20 in PBS and mounted. For Wnt5a antibody staining an antigenic retrieval protocol prior immunolabeling was applied. Slices were incubated in antigen retrieval solution (DAKO) for 10 min in a pressure cooker (115-118 °C and 70-80 kPa) and cooled down in tap water.

Cultured GCP were blocked with 10% NDS in PBS for 30 min at RT. Anti-Pax6 primary antibody (rabbit polyclonal anti-Pax6, BioLegend, 901301, 1:500) in 2% NDS in PBS was incubated at 4 °C over-night. Incubation with the secondary antibodies (Donkey polyclonal anti-rabbit Alexa Fluor-488, Invitrogen, A21206 and donkey polyclonal anti-rabbit Alexa Fluor-555, Invitrogen, A31572) together with DAPI was done during 1 h at room temperature.

For FISH, *Dab1 in situ* probes are described in ^10^, *Wnt5a* and *Fzd2* probes were generated following the same protocol described in ^10,42^. The primer sequences used for each probe are: *Wnt5A*-fw AGTCCCACTCCCAGGACC, *Wnt5A*-rw TCAGCTGGGCTAACACAAGA, *Fzd2*-fw ACCTAGCCTGCTCGCTATTTT and *Fzd2*-rv CCTCCAACCCAACCTATTTTTAC and were taken from Allen Brain Atlas (Probe RP_071127_03_A06 and Probe RP_080912_03_F07 respectively). FISH was performed as described ^10,42^.

### Quantitative real-time PCR

For quantitative real-time PCR, RNA from bEnd.3 cells or GCP culture was extracted using Trizol reagent and incubated with Turbo DNase (Invitrogen). RNA was reverse-transcribe into cDNA and performed the quantitative PCR assays as described in ^10^. TaqMan Gene Expression probes for mouse Wnt5a (Thermo Fischer, Mm00437347_m1), mouse mKi67 (Mm05911137_s1), mouse GADPH (Thermo Fischer, Mm 99999915_g1) and mouse ß2m (Thermo Fischer, Mm00437762_m1), the last two served as endogenous control for GCP and bEnd.3 cells respectively.

### Western blot

For Western blot analysis, cells were maintained in ice-cold RIPA buffer (50 mM Tris/HCl, 150 mM sodium chloride, 1% NP-40, 0.5% sodium deoxycholate, 0.1% SDS, 1 mM EDTA, 10 mM NaF, 1% complete protease inhibitor cocktail (complete EDTA-free proteinase inhibitor cocktail tablets), 1 mM sodium orthovanadate, and 0.5 mM dithiothreitol (DTT), pH 7.4) lysed. Protein content was determined using Pierce BCA Protein Assay Reagent according to the manufacturer’s instructions, and samples were separated by SDS-PAGE and transferred to nitrocellulose membranes. Membranes were blocked in TBS-T with skim milk powder (3%) and incubated with anti-Wnt5A (1:300, Abcam, ab229200) and anti-pan-cadherin (1:1000, Sigma, C1821) antibodies at 4°C overnight. Membranes were then incubated with HRP-conjugated secondary antibodies anti-rabbit HRP (1:2000, Jackson, 111-035-003) and anti-mouse HRP (1:2000, Jackson, 115-035-146) in a blocking solution for 1 hour at RT. HRP activity was detected using enhanced chemiluminescence (ECL) detection reagent and ImageQuant LAS 4000 system. Quantifications of Wnt5a signal were performed with Fiji and normalized with pan-cadherin signal as loading control.

### Electrophysiology

#### Slice preparation

3-4 weeks old mice were decapitated under deep isoflurane anesthesia. The cerebellum was removed and immersed into ice-cold (2-3°C) “slicing” solution containing (in mM): 240 sucrose, 5 KCl, 1.25 Na_2_HPO_4,_ 2 MgSO_4_, 1 CaCl_2_, 26 NaHCO_3_ and 10 D-Glucose, bubbled with 95% O_2_ and 5% CO_2_. Cerebellar vermis was isolated and parasagittal slices (220 μm thick) were cut using a vibratome (VT1200S, Leica). Slices were incubated for at least 1 h before recordings in ACSF containing (in mM): 125 NaCl, 2.5 KCl, 1.25 Na_2_HPO_4_, 1 MgSO_4_, 2 CaCl_2_, 26 NaHCO_3_ and 25 D-Glucose, bubbled with 95% O_2_ and 5% CO_2_, maintained at 32°C.

#### Electrophysiological apparatus and patch-clamp recordings

For electrophysiological recordings, slices were transferred to the recording chamber and perfused at 1.5 ml min^−1^ with ACSF in the presence of 100 μM picrotoxin unless stated otherwise, bubbled with 95% O_2_ and 5% CO_2,_ at room temperature (RT; 20–23 °C). Slices were visualized with an upright microscope equipped with a ×63, 0.9 NA water-immersion objective and differential interference contrast (DIC) optics (Scientifica, UK).

Patch pipettes were made from thick-walled borosilicate glass capillaries with filament (GB150F-8P, Science Product) by means of a Sutter P-1000 horizontal puller (Sutter Instruments, Novato, CA, USA). Recordings were obtained in patch-clamp cell-attached configuration or whole-cell configuration using an Axoclamp 200B amplifier (Molecular Devices, Union City, CA, USA).

For cell-attached recordings (10–80M*Ω* seal resistance), patch pipettes were filled with standard extracellular saline. Pipettes were held at 0 mV in the voltage-clamp mode. Consecutive 140 sec. current traces were filtered at 2 kHz and acquired at 50 kHz sampling rate. In these conditions, recorded spikes appear as biphasic current deflections. Recordings were judged to be stable when the shape and frequency of spikes was constant over time. Stable recordings were routinely obtained for as long as 20-30min.

For whole-cell recordings, after gigaseal formation the cell membrane was broken with small suctions and whole-cell established. Recordings were excluded if series or input resistances, assessed by -10mV voltage step and with membrane test command from Clampfit Software suite, varied by more than 20% over the time of the experiment. The liquid junction potential was not subtracted. The recording electrode was filled with an intracellular solution containing (in mM): 134 K-gluconate, 6 KCl, 4NaCl, 10 Hepes, 0.2 EGTA, 4 Mg-ATP, 0.3 Na3GTP, 14 Na2 phosphocreatine (pH 7.3, Osmolarity ∼290). For the extracellular stimulation a bipolar electrode with a patch pipette filled with saline solution was used. The pipette was placed on the slice in the molecular layer (∼ 1/3 of the distal part) lateral to the recorded Purkinje cell.

The input-output i.e. stimulus intensity- EPSC output was assessed with stimuli of increasing intensity from 0.5 to 10uA. The paired-pulse stimulation was obtained with two consecutive stimuli of 7-8 uA with 25ms interstimulus interval; the acquisition frequency was 0.33Hz. The long-term plasticity at the parallel fibers – Purkinje cell (PF-PC) synapse was assessed with a LTP protocol of stimulation at 1 Hz for 5 min in current-clamp mode. The responses were evoked with a pair of stimuli (7-8uA, 25 ms interstimulus interval) at 0.33 Hz, PCs were kept at -70mV. Plasticity was evaluated from the data points 5 min before tetanus and 20 min after. The PC intrinsic excitability was assessed in current-clamp mode by injecting 800ms step of currents from –100pA to 1000pA (100pA increments), and the current-frequency plot was obtained by the average spiking frequency over the step pulse; the action potential properties were calculated from the first action potential generated.

#### Data analysis

In cell-attached recordings, firing frequency (Inst. Freq.) of the tonic simple spikes was evaluated with the event-sorting function in pClamp 10 Software suite; the variability of the interspike interval (ISI) was calculated from the mean ISI and the ISI coefficient of variation (CV(ISI)) computed as standard deviation of interspike intervals (ISI)/ mean of ISI. Each Freq. and CV(ISI) value was computed from an event-sample recorded over a time window of 140 sec. In whole-cell recordings, the EPSC amplitude was measured and averaged over 100 events per data point.

### Imaging and data processing

All in vivo experiments included animals from at least two different litters and cell culture experiments were repeated at least three times. Images were taken using a laser scanning confocal spectral microscope of a 1024x1024-pixel size. For cerebellar quantifications, sagittal sections from the vermis were selected for image acquisition and only III, IV-V and VI folia were imaged for quantifications except for DiI labeling.

For vascular analyses, pH3, ki67, Pax6 in the ML, DiI and cell culture, maximal projections of the whole thickness of the section were selected; for Pax6 in the GCL, EdU, Wnt5a signal, PC coverage, ML length and cell soma roundness, single snapshots were selected. Quantifications were done with ImageJ/Fiji (version 2.16.0/1.54p). Number of branch points and cells were quantified with the cell counter plugin. Area and distance were obtained after drawing a region or distance of interest and measures were given directly by the program. DiI+ PF coherency was obtained with orientation J plugin ^28^. PC dendrite tracing was done until the tertiary dendrite with ImageJ/Fiji Neuroanatomy-SNT plugin. The Wnt5a expression pattern in the cerebellar cortex was analyzed by measuring fluorescence intensity along the EGL and ML+PCL using the ImageJ/Fiji Plot Profile tool. Integrated signal intensity of the different layers was analyzed using Python by calculating the mean across all values per genotype and litter, respectively. Numerical integration of the area under the averaged intensity profile curves was performed using the composite trapezoidal rule and Simpson’s rule, implemented via SciPy functions.

For active synapse analysis, images were exclusively taken from lobules/folia III and IV of sagittal cerebellar sections (thickness: 80 µm). For Vglut1/Homer colocalization analysis, images were acquired from proximal and distal regions (to PC somas) of the ML at 63x magnification with 4x digital zoom, comprising a z-stack of 2 µm with a 0.13 µm step size. For Vglut2/Homer colocalization analysis, images of the entire ML were acquired at 63x magnification with 1x zoom, using a z-stack of 10 µm and a 0.13 µm step size. Image stacks were subjected to deconvolution using Huygens Professional software. Vglut1+ and Homer+ puncta were analyzed using the IMARIS spot detection tool. Synaptic activity was determined using the MATLAB colocalization plugin. A maximum distance threshold of 0.2 µm between the centroids of pre- and postsynaptic spots was applied to define synaptic coupling, consistent with parameters reported in previous studies employing IMARIS-based synaptic input assignment ^43^.

Vglut2+ puncta quantifications were obtained by merging Calbindin and Vglut2 staining and binarizing the images with ImageJ/Fiji. Vglut2+ puncta colocalizing with Purkinje cell (PC) dendrites were manually counted in the ML using the multipoint tool. Coordinates of Vglut2+ puncta along the PC dendrites were distributed into 10 equal segments of the ML using Python. Vglut2+ Homer+ colocalization in contact with the main PC dendrite was defined as an active synapse and annotated with the ImageJ/Fiji cell counter plugin.

Brightness and contrast from the images were adjusted with Adobe Photoshop. Figures were prepared with Adobe Illustrator.

### Statistics

Data was obtained with measurements taken from distinct samples. Statistical analysis was performed using GraphPad Prism (version 8). First, normal distribution of the samples was tested. As samples followed a normal distribution, 2-tailed unpaired Student’s t-test was used. Statistical significance was designed as P < 0.05 (*), P < 0.01 (**) and P < 0.001 (***). All graph values indicate mean ± Standard Error of the Mean (SEM).

## Acknowledgements

We thank U. Bauer, P. Brendel, D. G. Krishnamoorthy, T. Belefkih, and D. Schmelzer for technical support, Dr. B. C. Kirchmaier for animal welfare documentation, and Auss Abbood for initial exploratory experiments. This work was supported by grants from European Research Council (ERC_AdG_Neurovessel_project 669742), the Deutsche Forschungsgemeinschaft (SFB1080 221828878, SFB1531 456687919, FOR2325 269353708, SFB1507 450648163, GRK2566 414985841, EXC2026-CPI 390649896 to A.A-P and SE 3181/1-1 to M.S.) and the Max Planck Fellow Program A.A-P.

## Author contributions

M.P. designed and performed the in vivo cerebellar experiments, including immunohistochemistry, fluorescent in situ hybridization, qPCR analyses, and quantitative morphometric analyses, with experimental assistance from R.H., J.R.N., F.S., N.A., C.L-C., and M.R.A., J.J. performed and analyzed with help from R.H. in vitro granule cell and Purkinje cell culture experiments and immunoblot analyses. A. D’E. designed, performed, and analyzed all electrophysiological recordings. S.S. carried out endothelial cell siRNA experiments and associated molecular analyses. A.A.-P., M.S., and M.P. conceived and designed the study. A.A.-P. supervised all stages of the project. M.P., M.S., and A.A.-P. wrote the manuscript. All authors discussed the results, interpreted the data, and provided input on the final manuscript.

---

**Fig. S1.**
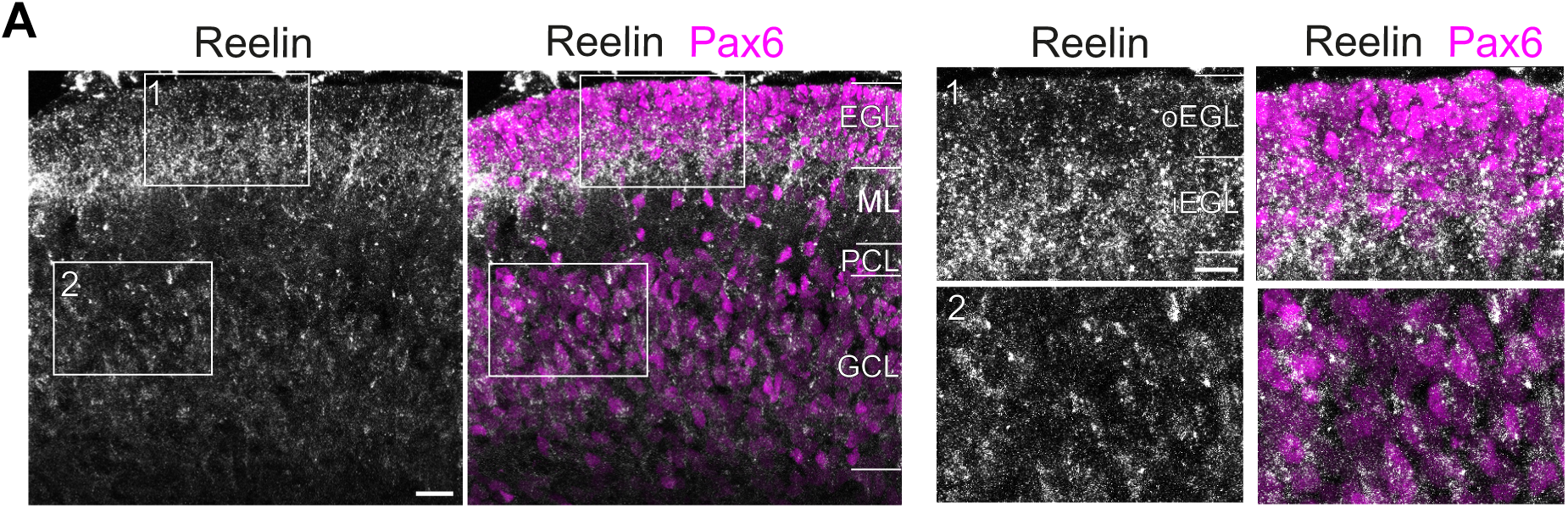
Reelin is expressed by postmitotic granule cells during postnatal cerebellar development. (**A**) Reelin immunostaining in cerebellar cortex at P8. Reelin is expressed by granule cells, detected with anti-Pax6 antibody. Higher magnification images on the right. Abbreviations: EGL = external granule layer, iEGL = inner EGL, ML = molecular layer, oEGL = outer EGL, PCL = Purkinje cell layer and GCL = granule cell layer. Scale bars: 20 µm (A), 10 µm (magnified region 1 and 2).

**Fig. S2.**
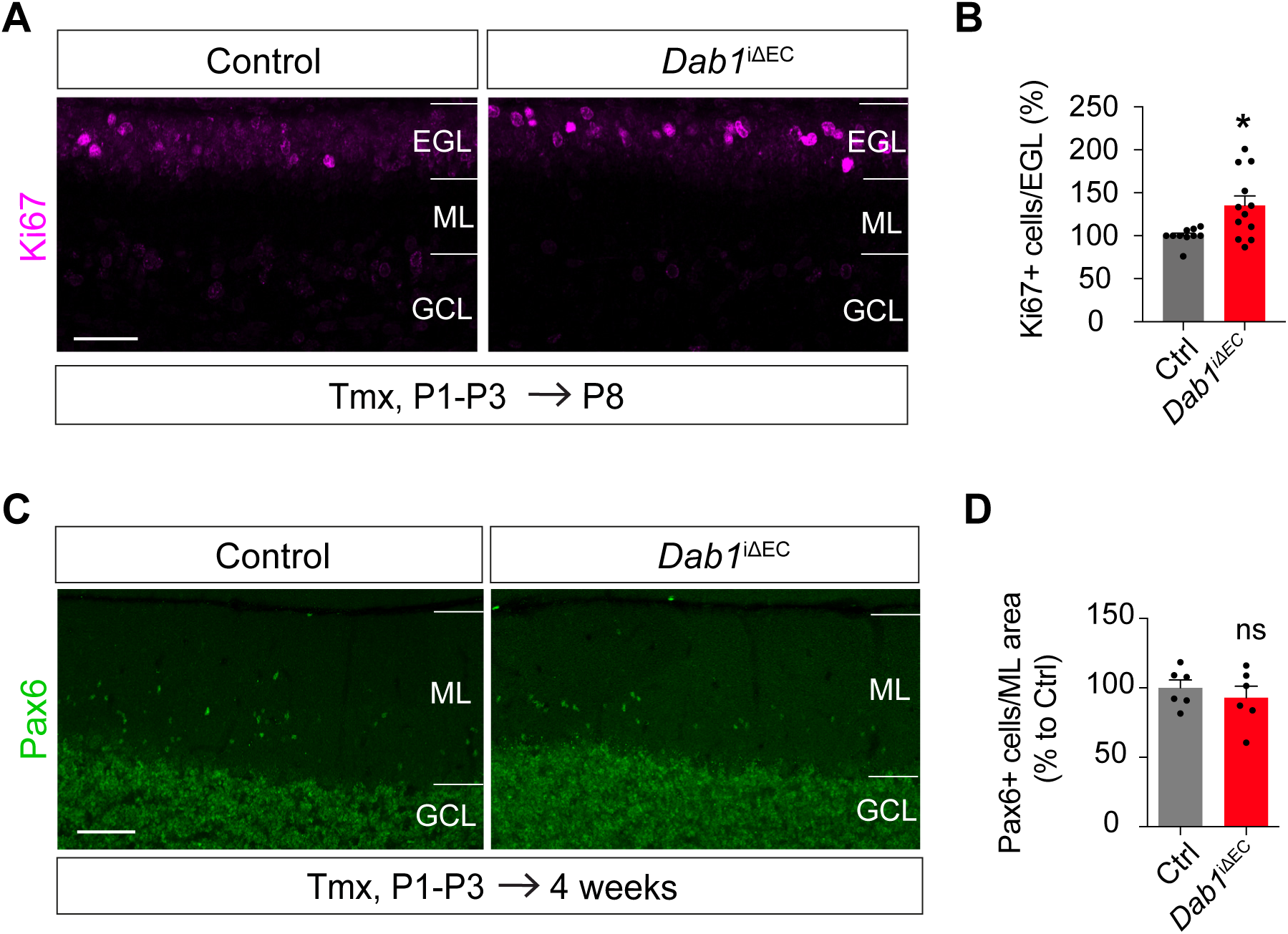
Endothelial Dab1 selectively regulates granule cell progenitor proliferation without altering migration. (**A**) anti-Ki67 immunostaining of *Dab1*^iΔEC^ and control cerebellar cortices at P8 after tamoxifen (Tmx) administration from P1 to P3. (**B**) Quantification of strongly labeled Ki67+ cells in (A) (n = 10-12 animals per genotype). (**C**) anti-Pax6 immunostaining in 4-week-old mice after Tmx administration from P1-P3. (**D**) Quantification of Pax6+ cells per area in the molecular layer in (C) (n = 6 animals per genotype). Abbreviations: EGL = external granule layer, GCL = granule cell layer and ML = molecular layer. Scale bars: A and C = 50 µm. Data are shown as mean ± SEM. \**P*<0.05, ns = non significant.

**Fig. S3.**
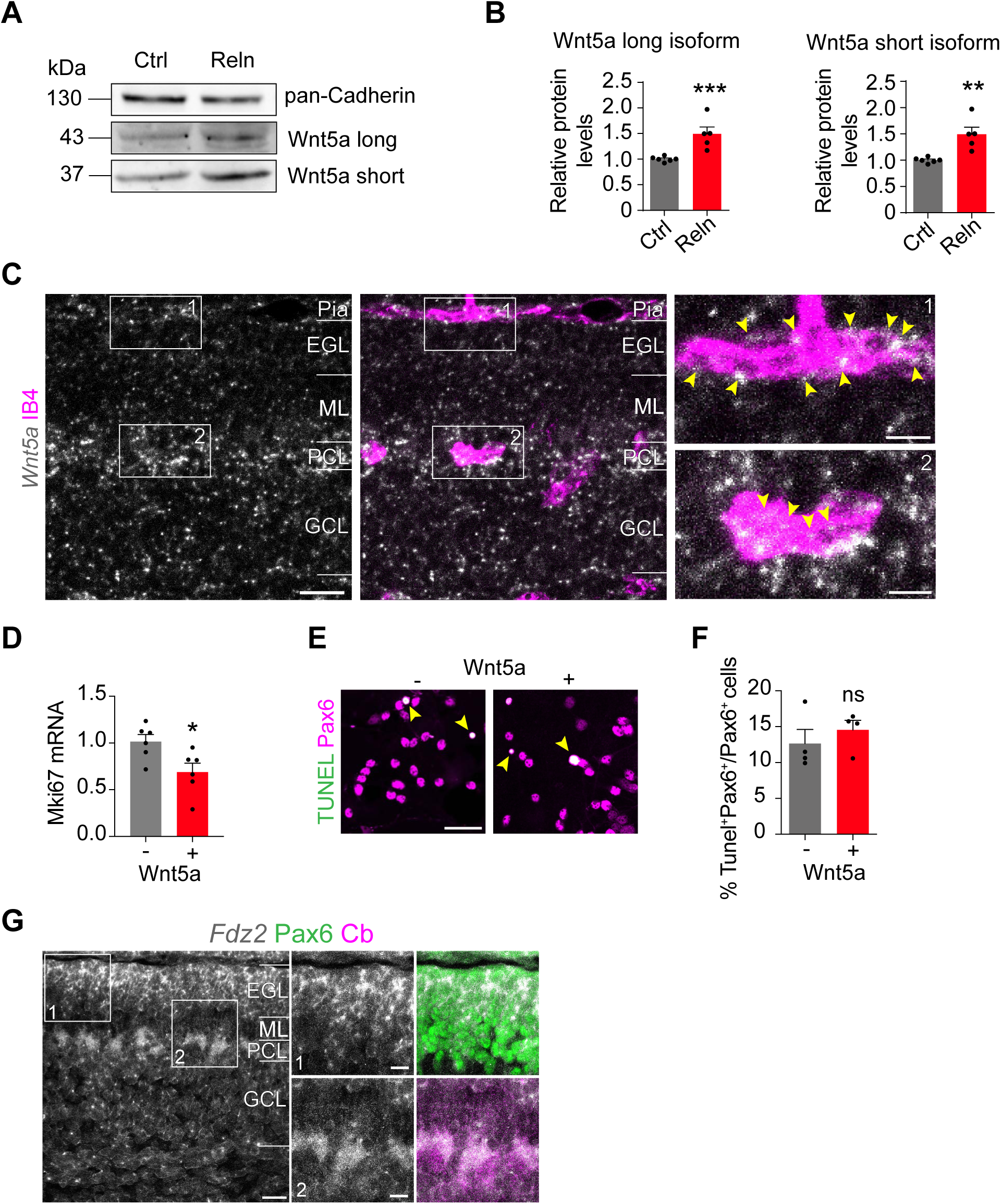
Endothelial-derived Wnt5a signals through Fzd2 to regulate granule cell progenitor proliferation. (**A**) Representative immunoblot showing enhanced Wnt5a protein expression in bEnd.3 cells after 48h stimulation with exogenous Reelin (Reln) protein. Pan-cadherin was used as a loading control. (**B**) Relative protein levels quantification of Wnt5a long isoform and short isoform versus pan-cadherin housekeeping protein in (A) (n = 3). (**C**) *Wnt5a* mRNA detected with fluorescence *in situ* hybridization in cerebellar cortex at P8. Blood vessels are labelled with isolectinB4 (IB4). Higher magnification images are shown on the right. Arrows indicate *Wnt5a* signal in a pial (1) and a cerebellar cortical vessel below the Purkinje cell layer (PCL) (2). (**D**) *Mki67* mRNA expression is increased in cerebellar granule cell progenitors (GCP) after 24 h stimulation with exogenous Wnt5a protein, assessed by quantitative PCR (n =6). (**E**) TUNEL and anti-Pax6 immunostaining of primary cell culture of GCP stimulated with exogenous Wnt5a protein. Arrows indicate apoptotic cells (TUNEL+ Pax6+). (**F**) Quantification of the proportion of apoptotic GCP in (E) (n =4). (**G**) *Fzd2* mRNA labelled by fluorescent in situ hybridization in cerebellar cortex at P8. Higher magnification images show granule cell progenitors immunolabeled with anti-Pax6 and Purkinje cells with anti-Calbindin (Cb) respectively. Abbreviations: EGL = external granule layer, GCL = granule cell layer, ML = molecular layer and PCL = Purkinje cell layer. Scale bars: C and E = 30 µm; G = 25 µm; squares in C and G =10 µm. Data are shown as mean ± SEM. \**P*<0.05, \*\**P*<0.01, \*\*\**P*<0.001, ns = non significant.

## References

1. D’Angelo, E., and Casali, S. (2012). Seeking a unified framework for cerebellar function and dysfunction: from circuit operations to cognition. Front Neural Circuits 6, 116. 10.3389/fncir.2012.00116.

2. Van Overwalle, F., Manto, M., Cattaneo, Z., Clausi, S., Ferrari, C., Gabrieli, J.D.E., Guell, X., Heleven, E., Lupo, M., Ma, Q., et al. (2020). Consensus Paper: Cerebellum and Social Cognition. Cerebellum 19, 833–868. 10.1007/s12311-020-01155-1.

3. Leto, K., Arancillo, M., Becker, E.B., Buffo, A., Chiang, C., Ding, B., Dobyns, W.B., Dusart, I., Haldipur, P., Hatten, M.E., et al. (2016). Consensus Paper: Cerebellar Development. Cerebellum 15, 789–828. 10.1007/s12311-015-0724-2.

4. Consalez, G.G., Goldowitz, D., Casoni, F., and Hawkes, R. (2020). Origins, Development, and Compartmentation of the Granule Cells of the Cerebellum. Front Neural Circuits 14, 611841. 10.3389/fncir.2020.611841.

5. Rahimi-Balaei, M., Bergen, H., Kong, J., and Marzban, H. (2018). Neuronal Migration During Development of the Cerebellum. Front Cell Neurosci 12, 484. 10.3389/fncel.2018.00484.

6. Cerminara, N.L., Lang, E.J., Sillitoe, R.V., and Apps, R. (2015). Redefining the cerebellar cortex as an assembly of non-uniform Purkinje cell microcircuits. Nat Rev Neurosci 16, 79–93. 10.1038/nrn3886.

7. Sillitoe, R.V., and Joyner, A.L. (2007). Morphology, molecular codes, and circuitry produce the three-dimensional complexity of the cerebellum. Annu Rev Cell Dev Biol 23, 549–577. 10.1146/annurev.cellbio.23.090506.123237.

8. Vogenstahl, J., Parrilla, M., Acker-Palmer, A., and Segarra, M. (2022). Vascular Regulation of Developmental Neurogenesis. Front Cell Dev Biol 10, 890852. 10.3389/fcell.2022.890852.

9. Di Marco, B., Crouch, E.E., Shah, B., Duman, C., Paredes, M.F., Ruiz de Almodovar, C., Huang, E.J., and Alfonso, J. (2020). Reciprocal Interaction between Vascular Filopodia and Neural Stem Cells Shapes Neurogenesis in the Ventral Telencephalon. Cell Rep 33, 108256. 10.1016/j.celrep.2020.108256.

10. Segarra, M., Aburto, M.R., Cop, F., Llao-Cid, C., Hartl, R., Damm, M., Bethani, I., Parrilla, M., Husainie, D., Schanzer, A., et al. (2018). Endothelial Dab1 signaling orchestrates neuro-glia-vessel communication in the central nervous system. Science 361. 10.1126/science.aao2861.

11. Li, S., Kumar, T.P., Joshee, S., Kirschstein, T., Subburaju, S., Khalili, J.S., Kloepper, J., Du, C., Elkhal, A., Szabo, G., et al. (2018). Endothelial cell-derived GABA signaling modulates neuronal migration and postnatal behavior. Cell Res 28, 221–248. 10.1038/cr.2017.135.

12. Kullmann, J.A., Trivedi, N., Howell, D., Laumonnerie, C., Nguyen, V., Banerjee, S.S., Stabley, D.R., Shirinifard, A., Rowitch, D.H., and Solecki, D.J. (2020). Oxygen Tension and the VHL-Hif1alpha Pathway Determine Onset of Neuronal Polarization and Cerebellar Germinal Zone Exit. Neuron 106, 607–623 e605. 10.1016/j.neuron.2020.02.025.

13. Howell, B.W., Hawkes, R., Soriano, P., and Cooper, J.A. (1997). Neuronal position in the developing brain is regulated by mouse disabled-1. Nature 389, 733–737. 10.1038/39607.

14. Mariani, J., Crepel, F., Mikoshiba, K., Changeux, J.P., and Sotelo, C. (1977). Anatomical, physiological and biochemical studies of the cerebellum from Reeler mutant mouse. Philos Trans R Soc Lond B Biol Sci 281, 1–28. 10.1098/rstb.1977.0121.

15. Goldowitz, D., Cushing, R.C., Laywell, E., D’Arcangelo, G., Sheldon, M., Sweet, H.O., Davisson, M., Steindler, D., and Curran, T. (1997). Cerebellar disorganization characteristic of reeler in scrambler mutant mice despite presence of reelin. J Neurosci 17, 8767–8777. 10.1523/JNEUROSCI.17-22-08767.1997.

16. Katsuyama, Y., and Terashima, T. (2009). Developmental anatomy of reeler mutant mouse. Dev Growth Differ 51, 271–286. 10.1111/j.1440-169X.2009.01102.x.

17. Sheldon, M., Rice, D.S., D’Arcangelo, G., Yoneshima, H., Nakajima, K., Mikoshiba, K., Howell, B.W., Cooper, J.A., Goldowitz, D., and Curran, T. (1997). Scrambler and yotari disrupt the disabled gene and produce a reeler-like phenotype in mice. Nature 389, 730–733. 10.1038/39601.

18. Borghesani, P.R., Peyrin, J.M., Klein, R., Rubin, J., Carter, A.R., Schwartz, P.M., Luster, A., Corfas, G., and Segal, R.A. (2002). BDNF stimulates migration of cerebellar granule cells. Development 129, 1435–1442. 10.1242/dev.129.6.1435.

19. Subashini, C., Dhanesh, S.B., Chen, C.M., Riya, P.A., Meera, V., Divya, T.S., Kuruvilla, R., Buttler, K., and James, J. (2017). Wnt5a is a crucial regulator of neurogenesis during cerebellum development. Sci Rep 7, 42523. 10.1038/srep42523.

20. Korn, C., Scholz, B., Hu, J., Srivastava, K., Wojtarowicz, J., Arnsperger, T., Adams, R.H., Boutros, M., Augustin, H.G., and Augustin, I. (2014). Endothelial cell-derived non-canonical Wnt ligands control vascular pruning in angiogenesis. Development 141, 1757–1766. 10.1242/dev.104422.

21. Masckauchan, T.N., Agalliu, D., Vorontchikhina, M., Ahn, A., Parmalee, N.L., Li, C.M., Khoo, A., Tycko, B., Brown, A.M., and Kitajewski, J. (2006). Wnt5a signaling induces proliferation and survival of endothelial cells in vitro and expression of MMP-1 and Tie-2. Mol Biol Cell 17, 5163–5172. 10.1091/mbc.e06-04-0320.

22. Sepp, M., Leiss, K., Murat, F., Okonechnikov, K., Joshi, P., Leushkin, E., Spanig, L., Mbengue, N., Schneider, C., Schmidt, J., et al. (2024). Cellular development and evolution of the mammalian cerebellum. Nature 625, 788–796. 10.1038/s41586-023-06884-x.

23. Ryu, Y.K., Collins, S.E., Ho, H.Y., Zhao, H., and Kuruvilla, R. (2013). An autocrine Wnt5a-Ror signaling loop mediates sympathetic target innervation. Dev Biol 377, 79–89. 10.1016/j.ydbio.2013.02.013.

24. Morris, R.J., Beech, J.N., Barber, P.C., and Raisman, G. (1985). Early stages of Purkinje cell maturation demonstrated by Thy-1 immunohistochemistry on postnatal rat cerebellum. J Neurocytol 14, 427–452. 10.1007/BF01217754.

25. Sotelo, C., Rossi, F. (2013). Purkinje Cell Migration and Differentiation. Handbook of the Cerebellum and Cerebellar Disorders. Springer, Dordrecht. 10.1007/978-94-007-1333-8_9.

26. Arancillo, M., White, J.J., Lin, T., Stay, T.L., and Sillitoe, R.V. (2015). In vivo analysis of Purkinje cell firing properties during postnatal mouse development. J Neurophysiol 113, 578–591. 10.1152/jn.00586.2014.

27. Cook, A.A., Fields, E., and Watt, A.J. (2021). Losing the Beat: Contribution of Purkinje Cell Firing Dysfunction to Disease, and Its Reversal. Neuroscience 462, 247–261. 10.1016/j.neuroscience.2020.06.008.

28. Fonck, E., Feigl, G.G., Fasel, J., Sage, D., Unser, M., Rufenacht, D.A., and Stergiopulos, N. (2009). Effect of aging on elastin functionality in human cerebral arteries. Stroke 40, 2552–2556. 10.1161/STROKEAHA.108.528091.

29. Ichikawa, R., Hashimoto, K., Miyazaki, T., Uchigashima, M., Yamasaki, M., Aiba, A., Kano, M., and Watanabe, M. (2016). Territories of heterologous inputs onto Purkinje cell dendrites are segregated by mGluR1-dependent parallel fiber synapse elimination. Proc Natl Acad Sci U S A 113, 2282–2287. 10.1073/pnas.1511513113.

30. Lin, H., Magrane, J., Clark, E.M., Halawani, S.M., Warren, N., Rattelle, A., and Lynch, D.R. (2017). Early VGLUT1-specific parallel fiber synaptic deficits and dysregulated cerebellar circuit in the KIKO mouse model of Friedreich ataxia. Dis Model Mech 10, 1529–1538. 10.1242/dmm.030049.

31. Goffinet, A.M. (1983). The embryonic development of the cerebellum in normal and reeler mutant mice. Anat Embryol (Berl) 168, 73–86. 10.1007/BF00305400.

32. Andersson, E.R., Salto, C., Villaescusa, J.C., Cajanek, L., Yang, S., Bryjova, L., Nagy, II, Vainio, S.J., Ramirez, C., Bryja, V., and Arenas, E. (2013). Wnt5a cooperates with canonical Wnts to generate midbrain dopaminergic neurons in vivo and in stem cells. Proc Natl Acad Sci U S A 110, E602–610. 10.1073/pnas.1208524110.

33. Arredondo, S.B., Guerrero, F.G., Herrera-Soto, A., Jensen-Flores, J., Bustamante, D.B., Onate-Ponce, A., Henny, P., Varas-Godoy, M., Inestrosa, N.C., and Varela-Nallar, L. (2020). Wnt5a promotes differentiation and development of adult-born neurons in the hippocampus by noncanonical Wnt signaling. Stem Cells 38, 422–436. 10.1002/stem.3121.

34. Bodmer, D., Levine-Wilkinson, S., Richmond, A., Hirsh, S., and Kuruvilla, R. (2009). Wnt5a mediates nerve growth factor-dependent axonal branching and growth in developing sympathetic neurons. J Neurosci 29, 7569–7581. 10.1523/JNEUROSCI.1445-09.2009.

35. Keeble, T.R., Halford, M.M., Seaman, C., Kee, N., Macheda, M., Anderson, R.B., Stacker, S.A., and Cooper, H.M. (2006). The Wnt receptor Ryk is required for Wnt5a-mediated axon guidance on the contralateral side of the corpus callosum. J Neurosci 26, 5840–5848. 10.1523/JNEUROSCI.1175-06.2006.

36. Kaiser, K., Jang, A., Kompanikova, P., Lun, M.P., Prochazka, J., Machon, O., Dani, N., Prochazkova, M., Laurent, B., Gyllborg, D., et al. (2021). MEIS-WNT5A axis regulates development of fourth ventricle choroid plexus. Development 148. 10.1242/dev.192054.

37. Yeung, J., Ha, T.J., Swanson, D.J., Choi, K., Tong, Y., and Goldowitz, D. (2014). Wls provides a new compartmental view of the rhombic lip in mouse cerebellar development. J Neurosci 34, 12527–12537. 10.1523/JNEUROSCI.1330-14.2014.

38. Kaiser, K., Gyllborg, D., Prochazka, J., Salasova, A., Kompanikova, P., Molina, F.L., Laguna-Goya, R., Radaszkiewicz, T., Harnos, J., Prochazkova, M., et al. (2019). WNT5A is transported via lipoprotein particles in the cerebrospinal fluid to regulate hindbrain morphogenesis. Nat Commun 10, 1498. 10.1038/s41467-019-09298-4.

39. Sotelo, C., and Dusart, I. (2009). Intrinsic versus extrinsic determinants during the development of Purkinje cell dendrites. Neuroscience 162, 589–600. 10.1016/j.neuroscience.2008.12.035.

40. Luck, R., Karakatsani, A., Shah, B., Schermann, G., Adler, H., Kupke, J., Tisch, N., Jeong, H.W., Back, M.K., Hetsch, F., et al. (2021). The angiopoietin-Tie2 pathway regulates Purkinje cell dendritic morphogenesis in a cell-autonomous manner. Cell Rep 36, 109522. 10.1016/j.celrep.2021.109522.

41. Lee, H.Y., Greene, L.A., Mason, C.A., and Manzini, M.C. (2009). Isolation and culture of post-natal mouse cerebellar granule neuron progenitor cells and neurons. J Vis Exp. 10.3791/990.

42. Parrilla, M., Chang, I., Degl’Innocenti, A., and Omura, M. (2016). Expression of homeobox genes in the mouse olfactory epithelium. J Comp Neurol 524, 2713–2739. 10.1002/cne.24051.

43. Kuljis, D.A., Park, E., Telmer, C.A., Lee, J., Ackerman, D.S., Bruchez, M.P., and Barth, A.L. (2019). Fluorescence-Based Quantitative Synapse Analysis for Cell Type-Specific Connectomics. eNeuro 6. 10.1523/ENEURO.0193-19.2019.

